# Transactivation of EGFR by LCK underlies an activity-defined dependency that predicts response to EGFR inhibition in cholangiocarcinoma

**DOI:** 10.64898/2026.11.11.765714

**Authors:** Enis H. Ozmert, Jennifer Tomlinson, Ryan D. Watkins, Dong-Gi Mun, Erik Jessen, Danielle M. Carlson, Amro M. Abdelrahman, Meina Cai, Brandon Wilbanks, Hendrien Kuipers, Nate W. Werneburg, Caitlin B. Conboy, Mark J. Truty, Gregory J. Gores, Akhilesh Pandey, Rory L. Smoot

**Author notes:** These authors contributed equally (Enis H. Ozmert, Jennifer Tomlinson).

## Abstract

Cholangiocarcinoma (CCA) is an aggressive biliary tract malignancy with limited molecularly guided treatment options. The Src-family kinase LCK is a therapeutic vulnerability in CCA, yet its canonical T-cell substrates are minimally expressed in CCA cells, leaving its downstream effectors undefined. Unbiased proximity labeling in HuCCT1 cells identified EGFR within the LCK signaling neighborhood. LCK depletion or inhibition reduced EGFR phosphorylation, whereas recombinant LCK increased EGFR phosphorylation in a cell-free assay, supporting a noncanonical role for LCK upstream of EGFR. Given the failure of EGFR-directed therapies in unselected CCA populations, we hypothesized that EGFR activity, rather than abundance or mutation status, identifies EGFR-dependent tumors. Among 28 patient-derived xenografts ranked by pretreatment substrate-inferred EGFR activity, the activity score correctly classified afatinib response in all six models tested. An analogous EGFR activity score stratified primary tumors in two human CCA cohorts. Together, these findings identify a therapeutically actionable LCK-EGFR signaling axis and nominate EGFR activity as a functional biomarker to guide tumor selection for EGFR-directed therapy in CCA, warranting prospective clinical validation.

**Highlights:**

- LCK transactivates EGFR in cholangiocarcinoma cells
- Recombinant LCK increases EGFR phosphorylation in a cell-free system
- EGFR activity, not abundance, predicted afatinib response in 6 of 6 PDX models
- An analogous EGFR activity score classifies tumors in two human CCA cohorts

## INTRODUCTION

Cholangiocarcinoma (CCA) is an aggressive biliary tract cancer with rising incidence and limited treatment options.^1^ Most patients present with advanced disease or recur after resection, requiring systemic therapy.^2^ Standard therapy consisting of gemcitabine and cisplatin combined with immune checkpoint blockade targeting the PD-1/PD-L1 has limited durability, while targeted agents against *FGFR2* fusions and *IDH1* mutations benefit only a minority of patients.^3,4^ These limitations underscore the need to identify additional therapeutic dependencies and strategies to select patients most likely to benefit from targeting them.

Lymphocyte-specific protein tyrosine kinase (LCK) is a non-receptor Src-family kinase best characterized in T-cell signaling.^5^ LCK is implicated in several solid tumors, including CCA, where it has been linked to chemoresistance, migration, and invasion.^5–9^ We previously identified LCK as a tumor-intrinsic dependency in CCA and showed that it phosphorylates YAP at Tyr357, promoting YAP nuclear retention and transcriptional activity independently of LATS signaling. Targeting LCK with small-molecule tool compounds or siRNA nanoparticle approaches demonstrated therapeutic efficacy in multiple preclinical CCA models.^10–12^

However, systemic LCK inhibition also suppresses T-cell activation, which limits LCK itself as a systemic drug target.^13,14^ Biliary epithelial cells also lack the T-cell receptor complex, ZAP70, and LAT through which LCK acts in T cells, so what LCK engages in these cells remains an open question.^5,13,14^ Identifying LCK effectors in CCA epithelial cells could therefore reveal alternative targets for disrupting tumor-intrinsic signaling without directly inhibiting LCK in T cells. To identify such effectors, we used unbiased proximity labeling to map the LCK signaling neighborhood in CCA cells and identified EGFR as a candidate.

Epidermal growth factor receptor (EGFR) is a strong biliary tract cancer dependency in CRISPR screens^15^ and is frequently overexpressed in CCA.^16^ EGFR-mediated feedback activation contributes to resistance to FGFR inhibitors in *FGFR2* fusion-positive CCA and to lenvatinib in hepatocellular carcinoma, with EGFR inhibition enhancing treatment responses in these settings.^17,18^ Nevertheless, EGFR-directed therapies have not demonstrated consistent clinical benefit in biliary tract cancer, either in unselected populations or in patients selected based on KRAS mutation status. ^19,20^ This disconnect raises the possibility that EGFR abundance alone does not identify tumors that depend on EGFR signaling. Evidence from other cancers suggests that receptor phosphorylation and pathway activity may be more informative markers of response than expression alone.^21^ In CCA, however, strategies to prospectively identify EGFR-dependent tumors remain lacking, and the mechanisms sustaining EGFR activity are incompletely defined.

Here, we show that the LCK proximity interactome varies with EGFR activity and that genetic dependency data independently link LCK to EGFR in biliary tract cancer cells. Phosphoproteomic reanalysis and a cell-free kinase assay support LCK as a noncanonical upstream activator of EGFR.^11^ We ranked 28 patient-derived xenograft (PDX) models by pretreatment substrate-inferred EGFR activity, with prospective classifications matching afatinib responses in all six treated models. Afatinib also demonstrated efficacy in two immunocompetent syngeneic CCA models. An analogous EGFR activity score stratified primary tumors in two independent human CCA cohorts and correlated significantly with LCK activity, supporting the relevance of this axis in human disease. ^22,23^ Together, these findings identify a noncanonical LCK–EGFR signaling axis and provide a preclinical rationale for selecting CCA tumors for EGFR-directed therapy by pathway activity.

## RESULTS

### Proximity labeling identifies EGFR in the LCK signaling neighborhood

In HuCCT1 CCA cells, the T-cell-restricted LCK substrates and adaptors CD247 (CD3ζ), CD3D, ZAP70, and LAT were minimally expressed, with mean normalized RNA-sequencing counts of 2.1, 4.1, 2.8, and 4.5, respectively, compared with 1,514 for EGFR (Table S1C). To map the LCK proximity interactome, we expressed LCK–TurboID–HA following doxycycline-induced depletion of endogenous LCK with sgLCK (Figure 1A).^24^ We expressed an LCK-TurboID-HA construct in cells lacking endogenous LCK after doxycycline-induced sgLCK, confining labeling to the fusion-defined LCK pool (Figure 1A). Immunoblotting confirmed loss of endogenous LCK and expression of the fusion protein (Figure 1B). Biotin labeling increased with dose in fusion-expressing cells and exceeded control-cell background (Figure S1A).

**Figure 1.**
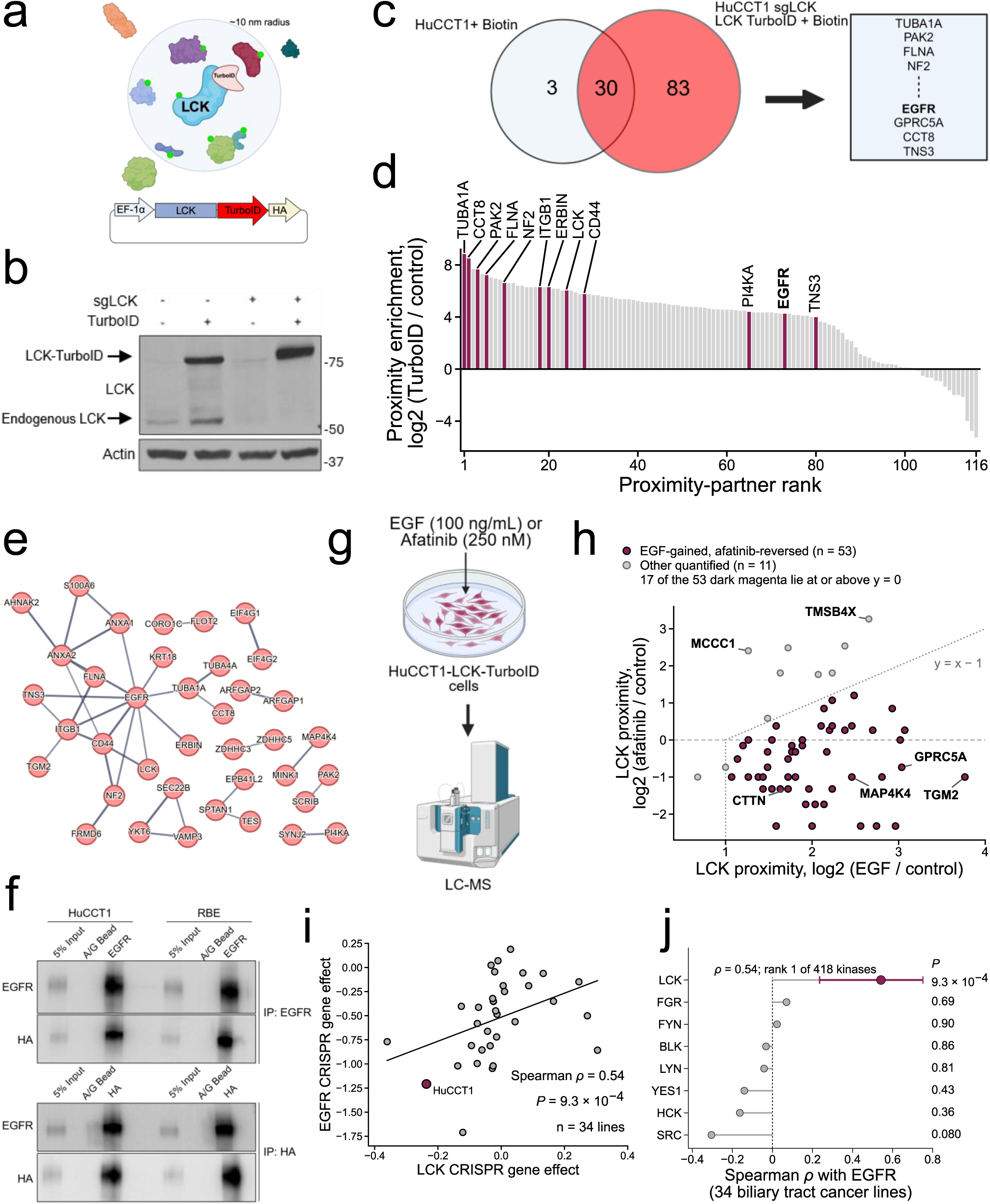
Proximity labeling identifies EGFR in the LCK signaling neighborhood in cholangiocarcinoma. (A) Schematic of the proximity-labeling strategy. (B) Immunoblot of HuCCT1 cells with and without doxycycline-induced sgLCK and with and without LCK-TurboID-HA expression. (C) Overlap of proteins recovered by streptavidin enrichment and mass spectrometry from biotin-treated control HuCCT1 cells (33 proteins) and biotin-treated HuCCT1-sgLCK/LCK-TurboID-HA cells (113 proteins). The boxed proteins at right, CCT8, EGFR, GPRC5A, TNS3, TUBA1A, PAK2, FLNA and NF2, were detected only in the fusion-expressing arm. The control is parental HuCCT1, and the mass-spectrometry experiment had one replicate per arm. (D) Proteins ranked by proximity enrichment, expressed as log2((LCK-TurboID intensity + 155,000) / (control intensity + 155,000)). All 116 proteins are ranked; the 12 labeled proteins, highlighted in dark magenta, are the two highest-ranked proteins (TUBA1A, CCT8), the kinases LCK, EGFR, PAK2 and PI4KA, and the adhesion-and cytoskeleton-associated proteins CD44, ITGB1, FLNA, NF2, ERBIN and TNS3. (E) STRING interaction network (v12; minimum combined score 0.7, lowest score displayed 0.706) of the 38 connected proteins among the 83 proteins detected only in LCK-TurboID cells, with 37 interactions. (F) Reciprocal co-immunoprecipitation under basal culture conditions: EGFR immunoprecipitation (top) and HA immunoprecipitation (bottom), each immunoblotted for EGFR and HA, in HuCCT1 and RBE cells expressing LCK-TurboID-HA. Input, 5% of lysate. Control lanes, A/G beads alone. One experiment is shown. (G) Schematic of the perturbation proximity-labeling experiment. (H) Log2 change in LCK proximity enrichment after afatinib versus EGF, each relative to control. Of 93 proteins quantified, 64 are plotted; 29 with an afatinib/control ratio of 0 are not. Dark magenta (53 proteins), log2 EGF/control above 1 and log2 afatinib/control more than 1 below log2 EGF/control; grey, the other 11. For 17 of the 53, the afatinib/control ratio is at or above 1. (I) Correlation between LCK and EGFR CRISPR gene effect (Chronos) across 34 biliary tract cancer cell lines in DepMap release 26Q1; HuCCT1 is highlighted. Spearman ρ and P value as shown; line, linear fit. (J) EGFR co-dependency for each Src-family kinase, expressed as the Spearman correlation of its gene effect with EGFR gene effect across the same 34 lines. LCK ranked first of 418 kinases surveyed by uncorrected correlation. P values, two-sided Spearman test, not adjusted; error bar on LCK, bootstrap 95% CI (1,000 resamples). See also Figure S1 and Table S1.

Anti-biotin enrichment of biotinylated peptides (BioSITe) and mass spectrometry identified 116 proteins, including 83 detected only in the fusion arm, among them LCK, CD44, ITGB1, FLNA, NF2, ERBIN, PAK2, and TUBA1A (Figure 1C; Table S1). EGFR was identified by a single biotinylated peptide detected only in the fusion arm (Figures 1C and 1D). STRING analysis placed EGFR in a connected module with LCK, ITGB1, CD44, NF2, and FLNA (Figure 1E).^25^ Reciprocal co-immunoprecipitation supported their association under basal conditions in HuCCT1 and RBE cells expressing LCK–TurboID–HA: EGFR immunoprecipitation recovered the fusion protein, and HA immunoprecipitation recovered EGFR, whereas bead-only controls recovered neither (Figure 1F). These findings place EGFR in the LCK signaling neighborhood and support their association in CCA cells.

To test whether EGFR activity alters the LCK proximity interactome, we treated LCK– TurboID–HA HuCCT1 cells with EGF (100 ng/mL) or afatinib (250 nM) before labeling and liquid chromatography–mass spectrometry (Figure 1G). Of 64 LCK-proximal proteins detected under both treatments, 53 showed increased proximity labeling with EGF and reduced labeling with afatinib relative to EGF, including CTTN, MAP4K4, GPRC5A, and TGM2 (Figure 1H; Table S1). Thus, the LCK proximity interactome varies with EGFR activity.

### EGFR is a lineage-enriched biliary dependency with LCK is its top ranked kinase co-dependency

In the Cancer Dependency Map (DepMap), biliary tract cancer cell lines showed stronger EGFR dependence than other cell lines pooled (one-sided Mann–Whitney P = 2.0 × 10⁻), with a similar pattern in ampulla of Vater lines (Figure S1B).^26^ Within the biliary lineage, EGFR dependence ranked below MYC, KRAS, and PIK3CA but above ERBB2, IGF1R, ERBB3, FGFR2, and IDH1 (Figure S1C).

Across 34 biliary tract cancer lines, LCK and EGFR Chronos gene-effect scores correlated positively (Spearman ρ = 0.54, P = 9.3 × 10⁻), with HuCCT1 among the co-dependent lines (Figure 1I).^27^ LCK was the strongest positive EGFR co-dependency among 418 kinases and ranked tenth among 17,786 genes. Other Src-family kinases showed little or no correlation (FGR, FYN, BLK, and LYN) or negative correlations (YES1, HCK, and SRC; Figure 1J). LCK gene-effect scores ranged from −0.36 to +0.30, indicating covariation in vulnerability rather than broad LCK essentiality (Figure S1C). These data provide independent genetic support for a functional LCK-EGFR signaling axis.

### LCK transactivates EGFR and supports proliferative signaling in cholangiocarcinoma

To test the functional significance of the LCK–EGFR association, we reanalyzed our published three-state SILAC tyrosine phosphoproteome comparing control cells with pharmacologic LCK inhibition by NTRC 0652-0 and CRISPR disruption of LCK (sgLCK). Across 144 quantified phosphotyrosine sites, the perturbations produced concordant changes. EGFR pY1172 fell to 0.01– and 0.09-fold of control with NTRC 0652-0 and sgLCK, respectively, while pY1197 fell to 0.48– and 0.26-fold (Figure 2A). The LCK/SFK pY394 activation-loop peptide, shared with FYN, YES1, and SRC, decreased to 0.11– and 0.22-fold, whereas total EGFR remained at 0.90– and 0.80-fold (Figure 2B). Occupancy-corrected data showed reduced phosphorylation across the EGFR C-terminal regulatory region, ERBB2, ERBB3, SHC1 Y427, and MEK1/2 and ERK1/2 activation sites (Figure S2A). During a 9-day doxycycline-inducible sgLCK time course, EGFR pY1092 progressively decreased while total EGFR remained stable (Figure 2C).

**Figure 2.**
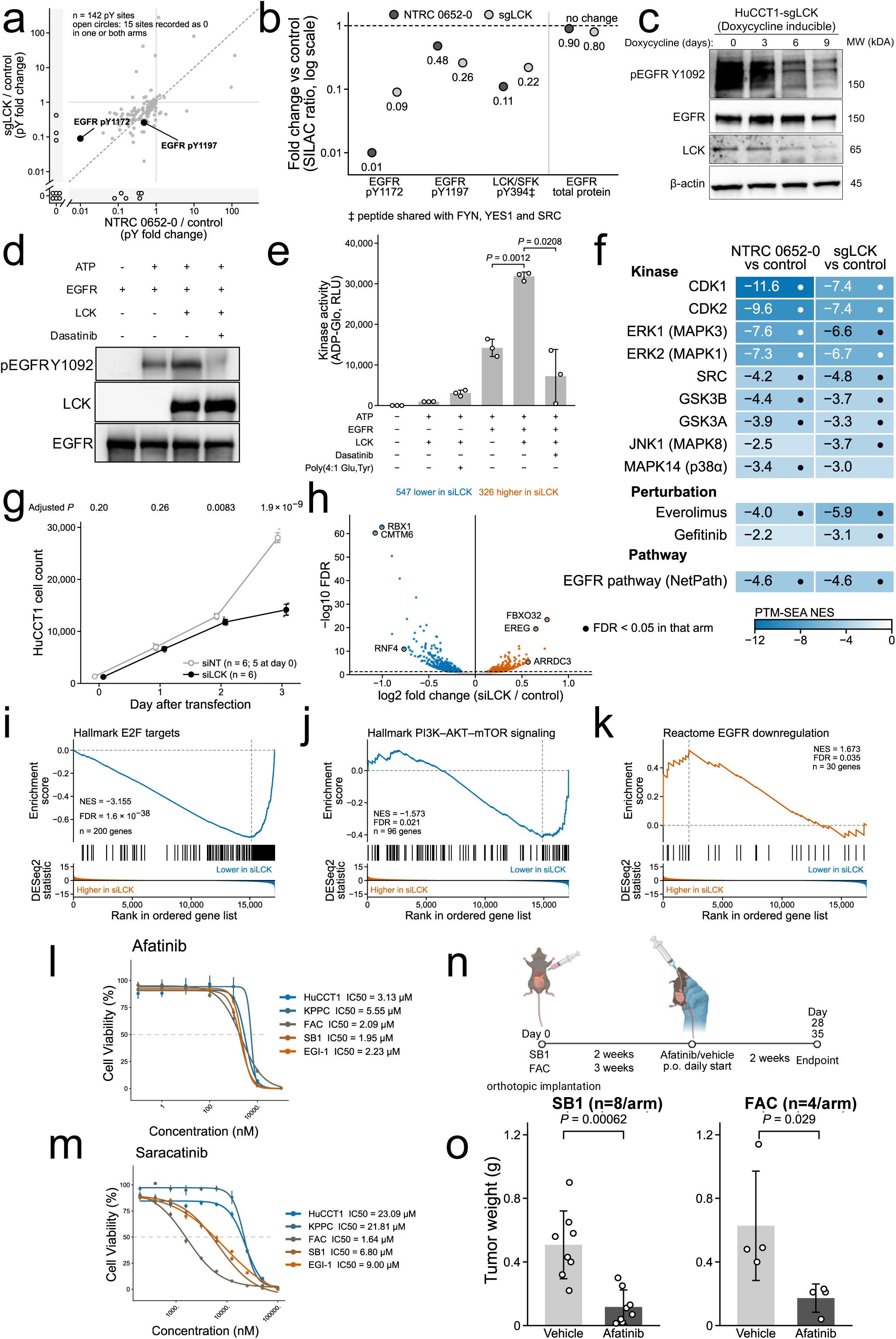
LCK transactivates EGFR and supports proliferative signaling in cholangiocarcinoma, and afatinib is efficacious in syngeneic orthotopic models. (A) Tyrosine phosphosite fold change in HuCCT1 cells following LCK inhibition with NTRC 0652-0 (x-axis) versus CRISPR disruption of LCK (y-axis). Of 144 quantified pY sites, the 142 with values in both arms are plotted. EGFR pY1172 and pY1197 are highlighted. Dashed line, identity. Values are plotted at their true positions on logarithmic axes; the 15 sites recorded as 0.00 in one or both arms are shown in the marked 0 strips beyond the axis breaks. Both arms come from one previously published three-state SILAC experiment (PXD026925). (B) Fold change versus control, on a logarithmic axis, for selected phosphotyrosine sites and total EGFR protein in the NTRC 0652-0 (dark grey) and sgLCK (light grey) arms of one SILAC experiment. LCK/SFK pY394 is a phosphopeptide shared with FYN, YES1 and SRC. Values below 1.0 indicate suppression relative to control. (C) Immunoblot of doxycycline-inducible HuCCT1-sgLCK cells over 9 days of doxycycline treatment, probed for pEGFR Y1092, total EGFR, LCK and β-actin. One experiment is shown. (D) Cell-free kinase reactions containing recombinant EGFR and recombinant LCK with the indicated components, immunoblotted for pEGFR Y1092, LCK and EGFR. Dasatinib, 500 nM. One experiment is shown. (E) ADP-Glo kinase activity of reactions assembled as in (D), with the components indicated and dasatinib at 500 nM, expressed as relative luminescence units. Bars, mean ± SD of three replicate reactions; individual replicates, open circles. Two-sided Welch’s t-test: EGFR + LCK versus EGFR alone, P = 0.0012; EGFR + LCK versus EGFR + LCK + dasatinib, P = 0.0208. P values compare replicate reactions. (F) PTM-SEA normalized enrichment scores for kinase, perturbation and pathway signatures (PTMsigDB v2.0.0) in the NTRC 0652-0 and sgLCK arms. Dots indicate FDR < 0.05 within that arm. (G) Proliferation of HuCCT1 cells transfected with siLCK or non-targeting siRNA (siNT), on days 0 through 3. Points, mean ± SD of 6 wells per condition from one experiment (5 siNT wells at day 0). Adjusted P values by Welch’s two-sample t-test with Benjamini-Hochberg correction across days. (H) Differential gene expression in HuCCT1 cells transfected with siLCK versus siNT, 72 h after transfection. 326 genes upregulated and 547 downregulated at FDR < 0.05. Selected genes are annotated. 3 biological replicates per condition. (I-K) Gene set enrichment analysis of the siLCK versus siNT comparison, ranked by the DESeq2 Wald statistic, for Hallmark E2F targets (I), Hallmark PI3K-AKT-mTOR signaling (J) and Reactome EGFR downregulation (K). NES and FDR as shown. (L and M) Dose-response curves and IC50 values for afatinib (L) and saracatinib (M) in human (HuCCT1, EGI-1) and murine (SB1, FAC, KPPC) CCA cell lines. Points, mean ± SD of 3 wells per condition from one experiment. (N) Schematic of the syngeneic orthotopic experiment: SB1 or FAC cells in the left liver lobe of C57BL/6 mice; afatinib or vehicle daily by oral gavage for 14 days, beginning 2 weeks (SB1) or 3 weeks (FAC) after implantation. (O) Tumor weight at endpoint for SB1 (n = 8 mice per arm) and FAC (n = 4 mice per arm; no mice excluded) orthotopic models treated with afatinib or vehicle. Bars, mean ± SD; points, individual animals. P values by two-sided exact Mann-Whitney U test: SB1, P = 0.00062; FAC, P = 0.029, the smallest value attainable at 4 versus 4. Hodges-Lehmann shift in tumor weight with afatinib, 0.37 g (SB1) and 0.28 g (FAC). See also Figure S2.

In a cell-free assay with recombinant EGFR and ATP, recombinant LCK increased EGFR pY1092, an effect blocked by dasatinib (500 nM; Figure 2D). ADP-Glo luminescence increased from 14,210 RLU with EGFR alone to 31,830 RLU with EGFR plus LCK (P = 0.0012), whereas LCK alone produced little signal. Dasatinib, which also inhibits EGFR, reduced luminescence to 7,246 RLU (P = 0.0208 versus EGFR plus LCK; Figure 2E). These findings support LCK as a noncanonical upstream activator of EGFR.

Post-translational modification signature enrichment analysis (PTM-SEA) showed negative enrichment of CDK1, CDK2, ERK1, ERK2, SRC, GSK3A, and GSK3B kinase signatures, the EGFR pathway signature, and the everolimus perturbation signature in both LCK perturbations (FDR < 0.05; Figure 2F).^28^ In HuCCT1 cells, siLCK lowered proliferation at day 2 (adjusted P = 0.0083) and day 3 (adjusted P = 1.9 × 10^−9^; Figure 2G) and impaired wound closure in HuCCT1 and SB1 cells (exact Mann-Whitney P = 0.0286 in each; Figures S2C-S2F). RNA sequencing identified 547 downregulated and 326 upregulated genes after siLCK (FDR < 0.05; Figure 2H). Hallmark E2F targets showed the strongest depletion (NES = −3.16, FDR = 1.6 × 10⁻³), alongside PI3K–AKT–mTOR depletion (NES = −1.57, FDR = 0.021; Figures 2I and 2J).^29^ Conversely, Reactome EGFR downregulation (NES = 1.67, FDR = 0.035; Figure 2K) and receptor tyrosine kinase signaling (FDR = 0.00063) were enriched. EGFR signaling and EGFR signaling in cancer showed nonsignificant trends (FDR = 0.080 and 0.088; Figure S2B). EREG was among the most induced transcripts, suggesting compensatory feedback (Figure 2H).

Across five human and murine CCA lines, afatinib reduced viability with IC₅₀ values ranging from 1.95 µM in SB1 to 5.55 µM in KPPC (Figure 2L). Saracatinib, a Src-family inhibitor with LCK activity, yielded IC₅₀ values of 1.64 µM in FAC to 23.09 µM in HuCCT1 (Figure 2M).^30^ Combination indices were below 1 in all five lines (0.16–0.99), at lower afatinib doses in EGI-1, SB1, and FAC and across the afatinib range, predominantly with low saracatinib, in HuCCT1 and KPPC (Figure S2G). SB1 and FAC, which had the lowest afatinib and saracatinib IC₅₀ values, respectively, are syngeneic to C57BL/6 mice. Following orthotopic liver implantation, immunocompetent mice received daily afatinib or vehicle for 14 days (Figure 2N). Afatinib reduced median endpoint tumor weight from 0.495 to 0.09 g in SB1 (n = 8 per arm; exact Mann–Whitney P = 0.00062) and from 0.485 to 0.21 g in FAC (n = 4 per arm; P = 0.029; Figure 2O).

### EGFR activity, rather than abundance, stratifies biliary tract cancer PDX models

We next asked whether EGFR activity, rather than abundance, identifies EGFR-dependent tumors. A median split on EGFR mRNA abundance did not stratify overall survival in either Fudan FU-iCCA (n = 201, log-rank P = 0.70) or Mayo CCA (n = 40, P = 0.89; Figure 3A).^22,23^

**Figure 3.**
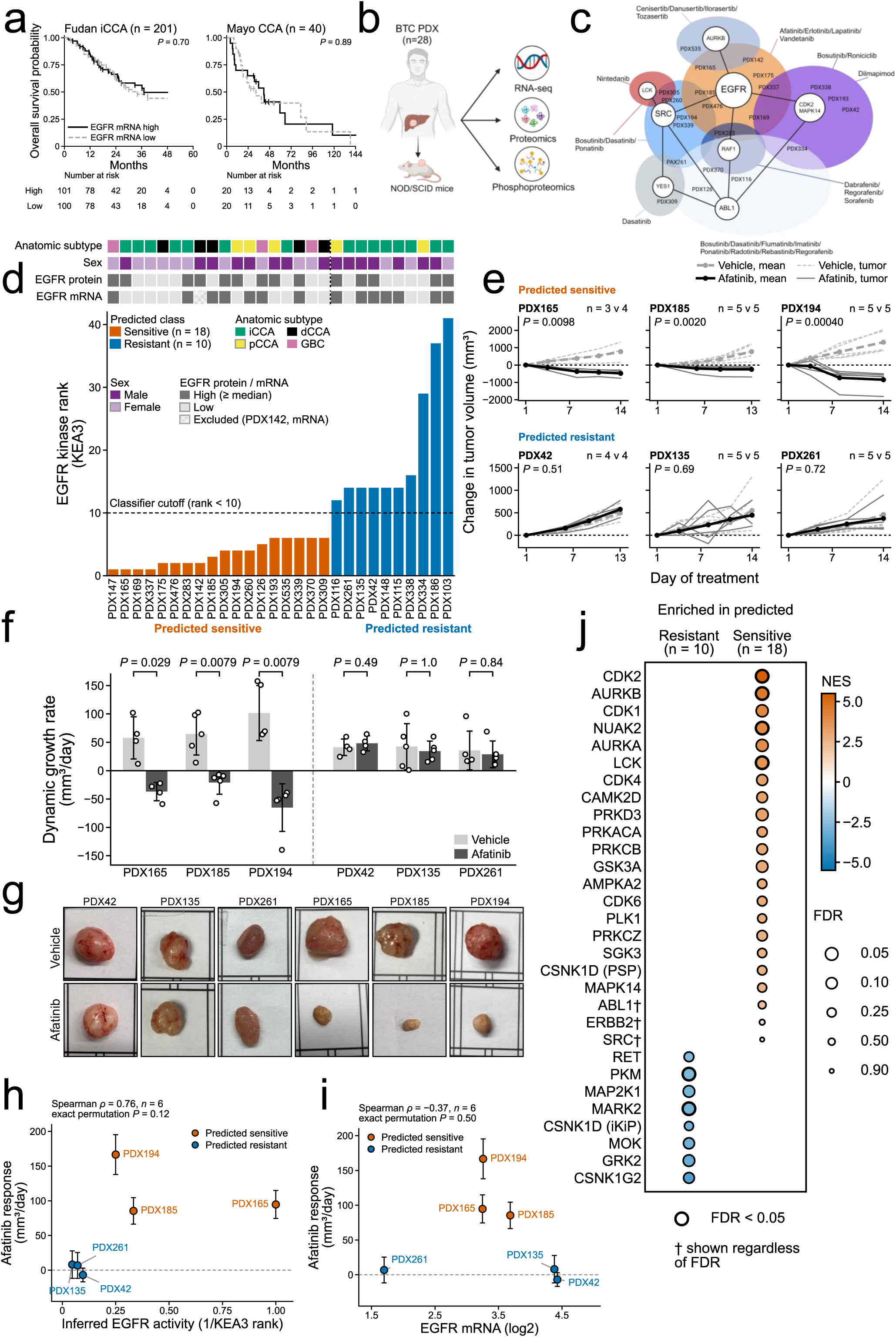
Inferred EGFR activity predicts afatinib response in PDX models. (A) Kaplan-Meier overall survival by EGFR mRNA expression in the Fudan FU-iCCA cohort (n = 201, left; log-rank P = 0.70) and an independent Mayo CCA cohort (n = 40, right; log-rank P = 0.89). Tumors were divided at the median of EGFR mRNA expression within each cohort. P values by log-rank test. Tick marks indicate censored patients; numbers at risk are shown below each plot. Excluded: 6 Fudan patients with no EGFR mRNA, 7 Fudan patients with no overall survival data, and 3 Mayo patients (2 who died within 90 days of surgery, 1 who had a palliative procedure only). (B) Schematic of the PDX profiling workflow. (C) Inferred kinase network centered on EGFR, annotated with clinically available inhibitors. (D) PDX models ranked by inferred EGFR kinase activity (KEA3 EGFR rank, 1 to 41; lower rank, higher activity). Annotation tracks show anatomic subtype, sex, EGFR protein level and EGFR mRNA level, each high (at or above the panel median) or low (below the median); PDX142 is shown as excluded from the mRNA track. Dashed line, classifier cutoff (rank below 10, predicted sensitive). Orange, predicted sensitive; blue, predicted resistant. (E) Change in tumor volume from baseline in six PDX models treated with afatinib (30 mg/kg daily by oral gavage; solid lines, dark grey to black) or vehicle (dashed lines, light to mid grey). Three predicted-sensitive models (top) and three predicted-resistant models (bottom). Thin lines, individual tumors; thick lines, arm means at each measured day. n per arm (vehicle/afatinib) is printed on the panel. (F) Dynamic growth rate for the same six models, calculated per mouse as the difference between the tumor volume on its last recorded day (day 14; day 13 for PDX185 and PDX42) and on day 1, divided by the number of days between the two measurements (mm^3^/day); negative values indicate regression. Bars, mean ± SD; points, individual mice. n as in (E), except PDX165 vehicle, n = 4 (the censored mouse is included, with its rate calculated to its last record, day 8). P values by two-sided exact Mann-Whitney U test, not adjusted across models. (G) Representative resected tumors from vehicle– and afatinib-treated mice for each model. (H) Afatinib response, expressed as the difference in mean dynamic growth rate (mm^3^/day) between vehicle and afatinib arms, versus inferred EGFR kinase activity score (1/KEA3 rank), across six PDX models. Error bars, propagated SEM of the two arms. PDX42, PDX135 and PDX261 share EGFR rank 14 and are offset horizontally for display. Spearman ρ = +0.76, exact permutation P = 0.12 (all 720 rank orderings), n = 6. (I) Afatinib response versus EGFR mRNA abundance across the six PDX models. Error bars as in (H). Spearman ρ = −0.37, exact permutation P = 0.50, n = 6. (J) Inferred kinase activity differing between predicted-resistant and predicted-sensitive PDX models by PTM-SEA (PTMsigDB v2.0.0). Dot color, normalized enrichment score; dot area, −log10 FDR. Black outline, FDR < 0.05; †, shown regardless of FDR (ABL1, ERBB2 and SRC). Orange, higher in predicted-sensitive; blue, higher in predicted-resistant. See also Figure S3 and Tables S2, S3 and S4.

We analyzed RNA-sequencing, proteomic, and phosphoproteomic profiles from 28 biliary tract cancer patient-derived xenograft (PDX) models and used KEA3 to infer sample-level kinase activity from phosphosite data (Figure 3B).^31^ EGFR mapped to a druggable kinase module containing LCK (Figure 3C). EGFR ranks ranged from 1 to 41 where 18 models ranked 1–6 and 10 ranked 12–41, prospectively designated predicted sensitive and predicted resistant, respectively (Figure 3D; Table S3). Class assignment was not associated with sex (P = 0.254), EGFR protein abundance (P = 0.944), EGFR transcript abundance (P = 0.309), or anatomic subtype (Fisher–Freeman–Halton exact P = 0.082; Figure 3D; Table S2; Figures S3E–S3H).

No measured EGFR phosphosite (T693, S991, T993, S1026, S1166, or Y1092) differed between classes (Figure S3A). A wider set of sites across EGFR, ERBB2, Src-family kinases, and associated signaling proteins also showed no separation (Figure S3B), and none of the 17,367 quantified sites, including 191 phosphotyrosine sites, differed significantly between classes (Figure S3C). Thus, substrate-inferred EGFR activity classified the models without significant differences in EGFR abundance or individual phosphosites.

### Substrate-inferred EGFR activity prospectively predicts afatinib response in vivo

We prospectively tested the classifier by treating six PDX models with afatinib: three predicted-sensitive models, PDX165, PDX185, and PDX194, and three predicted-resistant models, PDX42, PDX135, and PDX261. Each predicted-sensitive model regressed below baseline, whereas the predicted-resistant models kept growing and did not differ significantly from vehicle (Figure 3E). Growth-rate analysis confirmed negative growth rates in all three predicted-sensitive models and no significant drug effect in the three predicted-resistant models (Figure 3F), and representative resected tumors agreed with the longitudinal data (Figure 3G).

Predictions matched responses in all six models. Single-feature comparators matched responses in 4 of 6 models for EGFR protein abundance, 5 of 6 for batch-corrected EGFR pY1092, 2 of 6 for raw pY1092, and 3 of 6 for curated EGFR-substrate count (Table S4). Response correlations with inferred EGFR activity (Spearman ρ = 0.76, exact permutation P = 0.12; Figure 3H) and EGFR transcript abundance (ρ = −0.37, P = 0.50; Figure 3I) were not significant at n = 6. Only the EGFR activity classifier matched all six treatment responses.

### Inferred LCK activity is enriched in predicted-sensitive PDX models

To connect the EGFR classes to the upstream mechanism, we compared per-site phosphorylation differences between predicted-sensitive (n = 18) and predicted-resistant (n = 10) models by PTM-SEA using PTMsigDB v2.0.0. Inferred activity was higher in the predicted-sensitive class for LCK (NES = 3.68, FDR = 0.049), alongside CDK1, CDK2 (NES = 5.42, FDR < 0.05), CDK6, AURKA, AURKB (NES = 4.56, FDR < 0.05), PLK1, NUAK2 (NES = 4.02, FDR < 0.05), GSK3A, and several PKC isoforms (Figure 3J). Reference signatures for SRC (FDR = 0.898) and ERBB2 (FDR = 0.711) showed no class difference. Conversely, the predicted-resistant class had higher inferred activity of RET, MAP2K1, MARK2, PKM, MOK, GRK2, and the casein kinase 1 family, with MARK2 (NES = −3.00) and PKM (NES = −2.76) significant at FDR < 0.05.

Of 749 pathways tested, six were significant at FDR < 0.05: chromatin organization and epigenetic regulation of gene expression were enriched in the predicted-sensitive class, whereas keratinization, formation of the cornified envelope, cell-cell communication, and the RHO GTPase cycle were enriched in the predicted-resistant class (Figure S3D). These findings show that higher inferred LCK activity characterizes the EGFR activity-defined sensitive class.

### An analogous EGFR activity score stratifies primary human cholangiocarcinoma

Given that activity ranking classified PDX models with higher inferred LCK activity in the predicted sensitive class, we tested whether an analogous score stratifies primary human CCA. We used KEA3 with a z-score site-selection criterion distinct from the PDX criterion, standardized the score inside each cohort, and classified tumors scoring 0 or higher as predicted sensitive (STAR Methods). In the Fudan intrahepatic CCA (iCCA) cohort, the continuous score distribution defined 113 predicted sensitive and 101 predicted resistant tumors (Figure 4A, Table S3), with class unrelated to EGFR transcript abundance (P = 0.131) or sex (P = 0.891).^22^ In the Mayo cohort, the score defined 24 predicted sensitive and 19 predicted resistant tumors across intrahepatic, perihilar, and distal CCA, with class unrelated to EGFR transcript abundance (P = 0.913), anatomic subtype (P = 0.327), or sex (P = 0.369; Figure 4B).^23^

**Figure 4.**
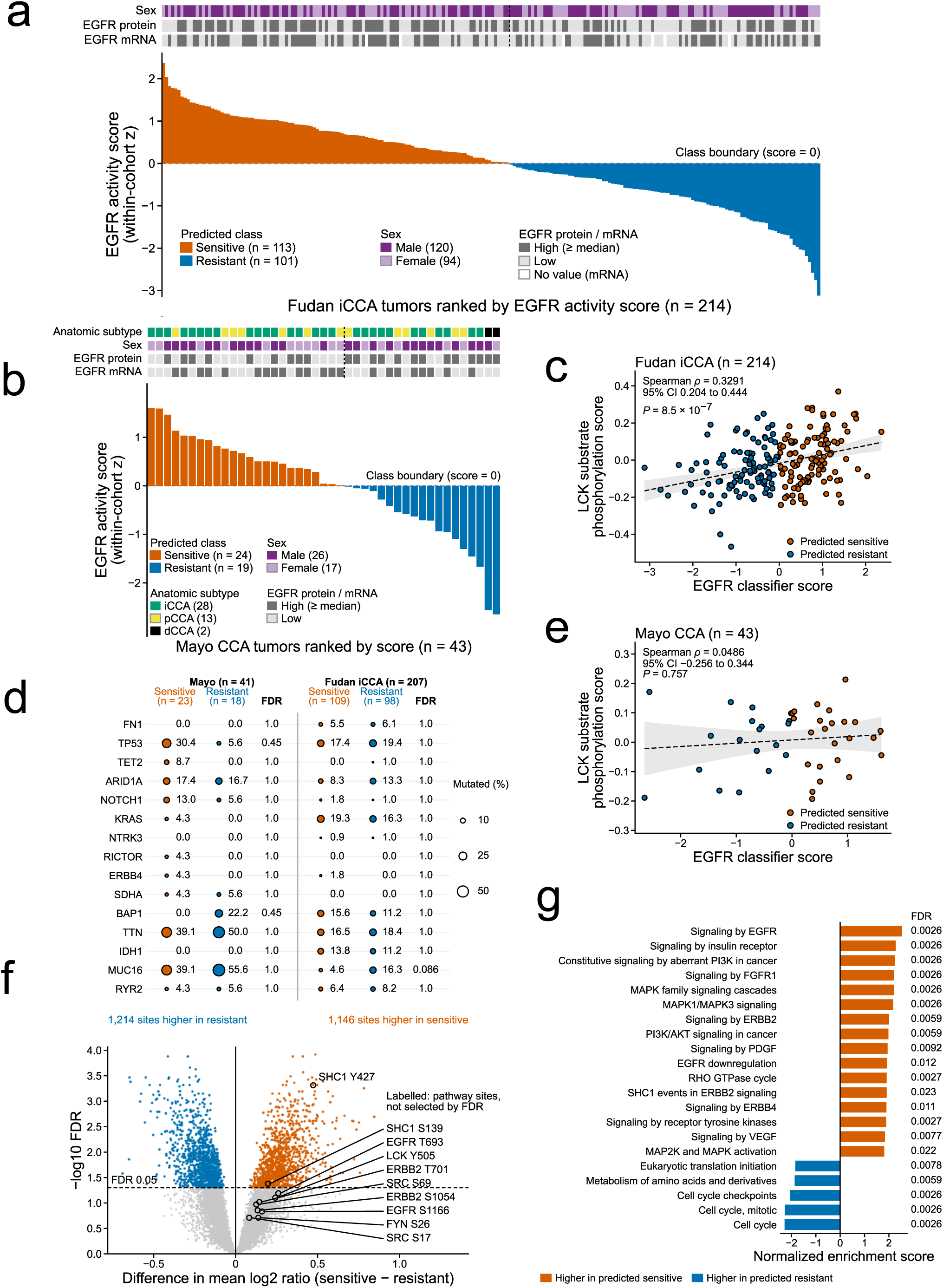
An analogous EGFR activity score stratifies primary human cholangiocarcinoma. (A) Tumors from the Fudan iCCA cohort (n = 214) ranked by EGFR kinase activity score standardized within the cohort. Tumors with a standardized score of 0 or higher (the within-cohort mean or above) are predicted sensitive (n = 113) and tumors below 0 predicted resistant (n = 101). Annotation tracks show EGFR protein level, EGFR mRNA level (no value for 6 tumors) and sex. For the protein and mRNA tracks only, high is at or above the within-cohort median of that measurement and low is below it (protein, 107 high and 107 low; mRNA, 104 and 104). The cohort is intrahepatic only, so no anatomic-subtype track is shown. Dashed line, class boundary. Orange, predicted sensitive; blue, predicted resistant. (B) Tumors from the Mayo CCA cohort (n = 43) ranked by EGFR kinase activity score, with classes defined as in (A): 24 predicted sensitive and 19 predicted resistant. Annotation tracks are as in (A), with anatomic subtype added. For the protein and mRNA tracks only, high is at or above the within-cohort median, so the median tumor is high, giving 22 high and 21 low for each track. (C) Frequency of coding-consequence variant calls in 15 genes in predicted-sensitive and predicted-resistant tumors of the Mayo (n = 41; 23 and 18) and Fudan (n = 207; 109 and 98) cohorts. Dot area indicates the mutated fraction; the adjacent column gives the Benjamini-Hochberg FDR across the 15 genes within each cohort (two-sided Fisher’s exact test per gene). The 15 genes are the most commonly mutated and CCA-relevant genes, and the genes not shown were also non-significant. IDH1 has no Mayo calls and cannot be tested there. Seven Fudan patients and 2 Mayo patients with no mutation analysis were excluded. (D) Inferred LCK activity (LCK-substrate phosphorylation score: unweighted mean of the matched OmniPath LCK-substrate phosphosites in the published log2-ratio table) versus EGFR classifier score in the Fudan cohort (n = 214). Spearman ρ = 0.3291 (95% CI 0.204 to 0.444), two-sided P = 8.5 × 10^−7^. Line, linear fit; shaded band, 95% confidence interval. Point color, predicted class. (E) Inferred LCK activity versus EGFR classifier score in the Mayo cohort (n = 43), quantified from TMT-ratio data. Spearman ρ = 0.0486 (95% CI −0.256 to 0.344), two-sided P = 0.757. Plotted as in (D). (F) Differential phosphosite abundance between predicted-sensitive (n = 113) and predicted-resistant (n = 101) tumors in the Fudan cohort, across 14,935 quantified phosphosites. Each point is one phosphosite. Orange, 1,146 sites higher in predicted-sensitive tumors; blue, 1,214 sites higher in predicted-resistant tumors (Benjamini-Hochberg FDR < 0.05); gray, not significant. Labeled sites include LCK pY505 and EGFR T693; phosphosite abundance reflects pathway engagement, including regulatory sites, and is not a direct measure of kinase activity. (G) Phosphoproteome pathway enrichment (fgsea; Reactome pathways) between predicted-sensitive (n = 113) and predicted-resistant (n = 101) tumors, ranked by normalized enrichment score. FDR is shown beside each bar. See also Tables S3 and S4.

By Mann-Whitney test, EGFR protein abundance was higher in predicted sensitive Fudan tumors (P = 0.0503) and predicted resistant Mayo tumors (P = 0.0454), although neither P value passed the Bonferroni reference thresholds (0.0167 and 0.0125). Overall survival did not differ by class in Fudan (log-rank P = 0.70) or Mayo (log-rank P = 0.29; Table S4). Across 15 selected genes (including TP53, ARID1A, KRAS, NOTCH1, BAP1, and IDH1), coding-variant frequency did not differ by class after Benjamini-Hochberg correction within Mayo (n = 41, minimum FDR 0.446) or Fudan (n = 207, minimum FDR 0.086), indicating that the classes do not recapitulate a recurrent genomic subtype among these genes (Figure 4C).

In the 214 Fudan tumors, inferred LCK activity (phosphorylation of LCK substrates) correlated positively with the EGFR classifier score (Spearman ρ = 0.3291, 95% CI 0.204 to 0.444, P = 8.5 × 10^−7^; Figure 4D). We adjusted for T-cell content (scored from each tumor’s own RNA-seq) in the 208 tumors with RNA-seq, and the association remained intact (partial Spearman ρ = 0.358, 95% CI 0.233 to 0.472, P = 1.2 × 10^−7^; unadjusted ρ = 0.337; Table S4). In the Mayo cohort (n = 43; 80% power for a correlation of 0.416 or larger), ρ was 0.0486 (95% CI −0.256 to 0.344, P = 0.757; Figure 4E) and the adjusted correlation was near zero (Table S4). Although the Mayo interval contained the Fudan estimate (difference between the two estimates P = 0.089), the association between inferred LCK activity and the EGFR score is supported in the larger Fudan cohort, holds after adjustment for T-cell content, and remains unconfirmed across independent patient datasets.

Differential phosphosite analysis of 14,935 quantified sites in the Fudan cohort identified 1,146 sites higher in predicted sensitive and 1,214 higher in predicted resistant tumors (FDR < 0.05; Figure 4F). Phosphoproteome pathway analysis showed enrichment of receptor tyrosine kinase signaling (EGFR, ERBB2, ERBB4, FGFR1, PDGF, VEGF, insulin receptor) plus downstream MAPK and PI3K-AKT in predicted sensitive tumors (FDR < 0.025), whereas predicted resistant tumors were enriched for cell cycle and translation (FDR < 0.009; Figure 4G).

Across the PDX models and both cohorts, the EGFR activity classes are not explained by EGFR protein or transcript abundance, indicating that EGFR dependency tracks the signaling state of the tumor, not receptor expression.

## DISCUSSION AND CONCLUSION

This study identifies a noncanonical LCK-EGFR signaling axis in cholangiocarcinoma (CCA) that connects an upstream Src-family kinase vulnerability to an activity-defined dependency on EGFR. Best known as a T-cell kinase and previously characterized in CCA as a YAP kinase, LCK also activates EGFR, a strong biliary tract cancer dependency.^10,11^ Specifically, the major findings are: (1) LCK is a noncanonical kinase that transactivates EGFR in CCA cells, where unbiased proximity labeling places EGFR in the LCK proximity interactome, LCK loss or pharmacologic inhibition lowers EGFR tyrosine phosphorylation, recombinant LCK increases EGFR phosphorylation in a cell-free assay, and the LCK proximity interactome follows EGFR activity as proteins gain proximity to LCK with EGF treatment and lose it with afatinib; (2) EGFR activity inferred from substrate phosphorylation, and not EGFR transcript or protein abundance, matched the afatinib response class of the treated patient-derived xenograft (PDX) models; and (3) an analogous EGFR activity score splits primary human CCAs in the Fudan and Mayo cohorts into activity-defined classes that are not explained by EGFR transcript abundance, anatomic subtype, or mutations in commonly altered genes. Our results therefore support the view that EGFR dependency in CCA follows the signaling state of the tumor, and LCK transactivation is one source of EGFR activity.

Prior studies of EGFR regulation by Src-family kinases have centered on other family members. c-Src associates with EGFR and regulates its phosphorylation at Tyr845 and Tyr1101.^32^ In rat hepatocytes, the Src-family kinase Yes associates with EGFR and Yes knockdown blunts EGFR activation under hyperosmotic stress, whereas the functional relevance of Lck activation there was not determined.^33^ Neither report tests LCK as a transactivator of EGFR in CCA cells. Our work therefore adds LCK, a lymphoid Src-family kinase, to the kinases known to regulate EGFR. Consistent with a distinct role for LCK in biliary epithelium, among the Src-family kinases examined in biliary tract cancer cells, only LCK has a strong positive EGFR co-dependency (ρ = 0.54; the other Src-family kinases ranged from −0.31 to +0.07). However, EGFR is the stronger genetic dependency and LCK has a near-neutral average CRISPR gene effect, suggesting that LCK may be a context-dependent activating input to EGFR, not a uniform survival requirement.

EGFR offers a clinically tractable downstream target, whereas systemic LCK inhibition suppresses proximal T-cell receptor signaling.^13,14^ Afatinib irreversibly inhibits EGFR, ERBB2, and ERBB4, blocking auto– and transphosphorylation within ErbB dimers and consequently signaling across the family.^34^ Its antitumor activity may therefore reflect an EGFR-centered ErbB dependency rather than dependence on EGFR alone. This interpretation connects LCK-mediated regulation to receptor-network inhibition without attributing afatinib efficacy exclusively to EGFR.

Despite experimental evidence of EGFR dependence, clinical trials have shown limited benefit.^15^ In a randomized phase 3 trial of erlotinib with gemcitabine and oxaliplatin, neither progression-free nor overall survival improved across the full study population.^35^ Trials of cetuximab and panitumumab with chemotherapy showed no consistent survival benefit, wild-type KRAS selection included.^19,20,36^ In a phase I study of afatinib with gemcitabine and cisplatin in nine patients with advanced CCA, nearly all with EGFR-overexpressing tumors, the authors found no survival benefit, and recruitment stopped because afatinib lacked efficacy in other indications.^37^ None of these studies prospectively identified tumors with active EGFR signaling; instead, they enrolled unselected patients or selected by EGFR abundance or KRAS status. These negative results show that enrolling unselected patients, or selecting by receptor abundance or KRAS status, fails as a strategy, and they do not establish EGFR as dispensable across CCA.

In contrast, our classifier, applied prospectively in six independently treated PDX models, separated afatinib-responsive from non-responsive tumors, which shows in preclinical models that signaling activity marks a therapeutically relevant subset. The predictive signal is spread across the phosphoproteome, as EGFR transcript or protein abundance and single receptor– or adaptor-associated phosphosites do not reproduce the classification into predicted sensitive and predicted resistant models. Predicted resistant tumors also have higher inferred activity of alternative kinases and enrichment of keratinization, cornified envelope, and cell-cell communication programs, suggesting that resistance goes with an alternative signaling state, not only absent EGFR activity. Conversely, inferred LCK activity is higher in the predicted sensitive class of PDX models than in the predicted resistant class, connecting upstream LCK signaling to the responsive state. Prospective phosphoproteomic profiling in precision-oncology programs has begun to link kinase activity scores to treatment response in other tumor types, including a phosphoproteomic biomarker distinguishing EGFR-inhibitor response in chordoma^38^; our data extend that logic to biliary tract cancer and identify an upstream kinase that contributes to EGFR activity.

An analogous, cohort-standardized score using a different site-selection criterion stratified primary tumors in the Fudan and Mayo cohorts. The classes spanned anatomic subtypes and were not significantly associated with coding variants in 15 selected genes, suggesting that functional pathway assessment could complement genomic selection. Although tumor-infiltrating T cells also express LCK, in the Fudan cohort LCK-substrate phosphorylation correlates with the EGFR score and survived adjustment for T-cell content, indicating a tissue-level relation that sits alongside the cell-line mechanism.^5^ These findings establish that LCK transactivation is a source of EGFR activity in CCA. Together, the results provide a mechanistic and preclinical rationale for prospective evaluation of activity-guided EGFR-directed therapy in CCA.

## RESOURCE AVAILABILITY

### Lead contact

Further information and requests for resources and reagents should be directed to and will be fulfilled by the lead contact, Rory L. Smoot.

### Materials availability

The HuCCT1-sgLCK/LCK-TurboID-HA and RBE LCK-TurboID-HA cell lines generated in this study are available from the lead contact with a completed materials transfer agreement.

### Data and code availability

The proximity-labeling mass-spectrometry proteomic data will be deposited in the ProteomeXchange Consortium through the PRIDE partner archive. The siLCK RNA-sequencing data from HuCCT1 cells will be deposited in the Gene Expression Omnibus. Both datasets will be released with reviewer access and made public on acceptance. The SILAC phosphoproteomic data analyzed in Figure 2 were reported previously and are available through PRIDE under accession PXD026925.^11^ The PDX and Mayo patient proteomic and phosphoproteomic data were previously reported (PRIDE PXD059245; SRA PRJNA1346066). The Fudan cohort data were previously reported and are available from the National Omics Data Encyclopedia under accession OEP001105.^22,23^ DepMap release 26Q1 is publicly available from the DepMap portal (https://depmap.org/portal/). Per-model PDX EGFR ranks and classes and per-patient cohort scores and classes are provided in Table S3. Any additional information required to reanalyze the data reported in this paper is available from the lead contact upon request.

## ACKNOWLEDGMENTS

We thank all the members of the GI Basic Research Program on Guggenheim Floor 17 at the Mayo Clinic, Rochester, MN.

## AUTHOR CONTRIBUTIONS

E.H.O., J.T. and R.L.S. conceived the study and its design. R.L.S. supervised the research. E.H.O., J.T., R.D.W., M.C., B.W., H.K., C.B.C. and N.W.W. contributed to in vitro studies. E.H.O., J.T., D.M.C. and A.M.A. performed in vivo studies, animal handling and tissue collection. M.J.T. contributed the PDX tissue repository. D.-G.M. and E.J. contributed to bioinformatic analysis. E.H.O. drafted the original manuscript. R.L.S., A.P. and G.J.G. contributed critical revisions of the manuscript. All authors reviewed the final manuscript. Funding was obtained by R.L.S. and G.J.G.

## DECLARATION OF GENERATIVE AI AND AI-ASSISTED TECHNOLOGIES IN THE MANUSCRIPT PREPARATION PROCESS

During the preparation of this work the authors used Claude (Anthropic) to revise manuscript text from the authors’ data and analyses, to write code that generated the figures directly from the analyzed data tables, to run and cross-check computational analyses, and to verify citations and numerical values against source records. GPT (OpenAI) and Gemini (Google) models were used only for automated quality review of the figures. After using these tools, the authors reviewed and edited the content as needed and take full responsibility for the content of the publication.

## SUPPLEMENTAL INFORMATION TITLES AND LEGENDS

**Figure S1.**
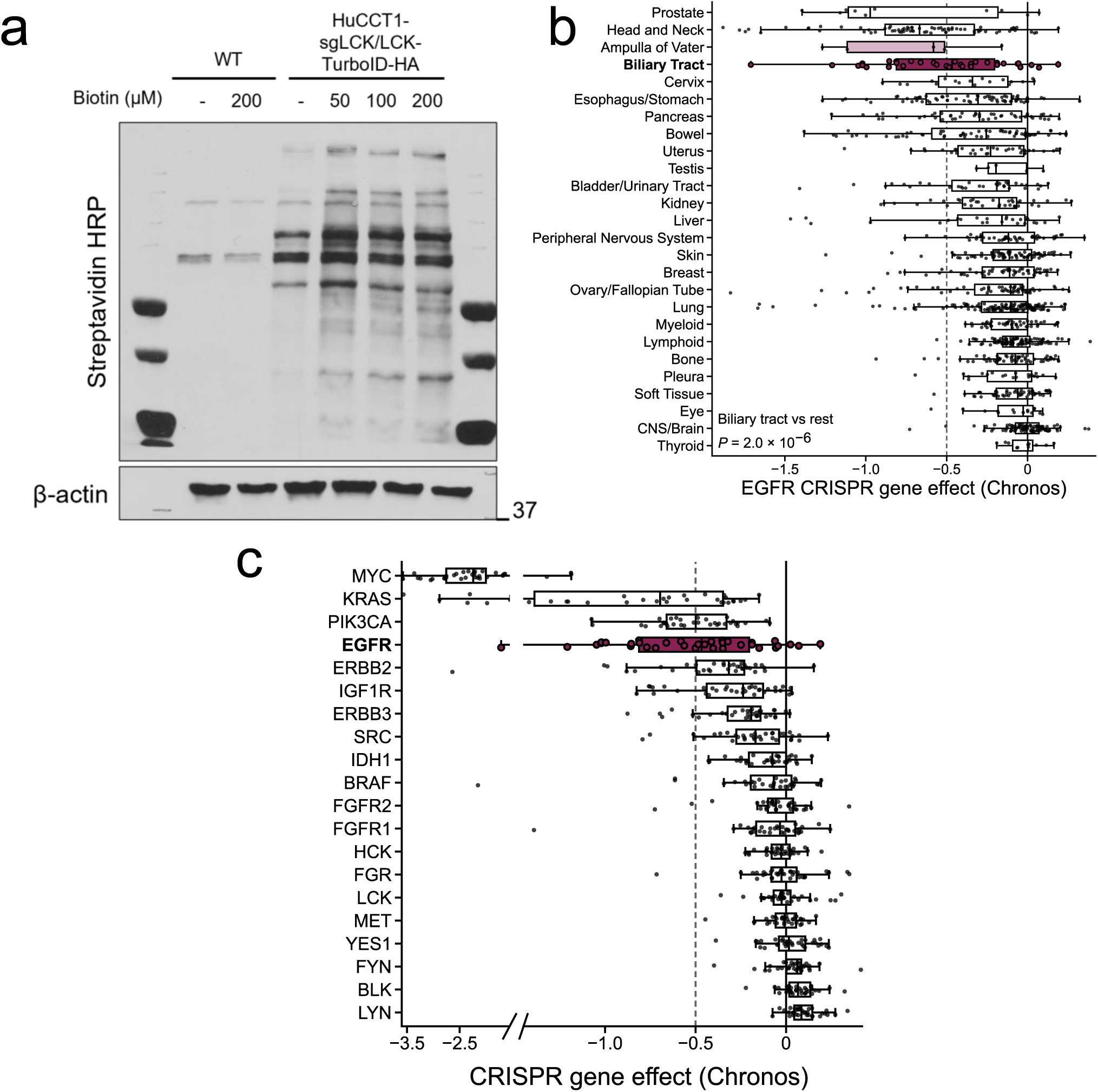
Biotin labeling optimization and EGFR dependency across cancer lineages, related to Figure 1. (A) Streptavidin-HRP immunoblot of control HuCCT1 and HuCCT1-sgLCK/LCK-TurboID-HA cells treated with the indicated biotin concentrations, with β-actin loading control. The control is parental HuCCT1. One experiment is shown. Blot images were not adjusted for brightness or contrast. (B) EGFR CRISPR gene effect (Chronos) across cancer lineages in DepMap release 26Q1. Biliary tract and ampulla of Vater lineages are highlighted. Boxes show median and interquartile range; whiskers extend to 1.5 times the interquartile range; points are individual cell lines; lineages with at least 5 lines are shown; the dashed line marks the essentiality threshold (−0.5). Biliary tract versus all other lineages, P = 2.0 × 10^−6^ by one-sided Mann-Whitney U test (two-sided P = 3.9 × 10^−6^). (C) CRISPR gene effect (Chronos) for selected cholangiocarcinoma-relevant genes across biliary tract cancer cell lines. EGFR is highlighted. Dashed line indicates the essentiality threshold.

**Figure S2.**
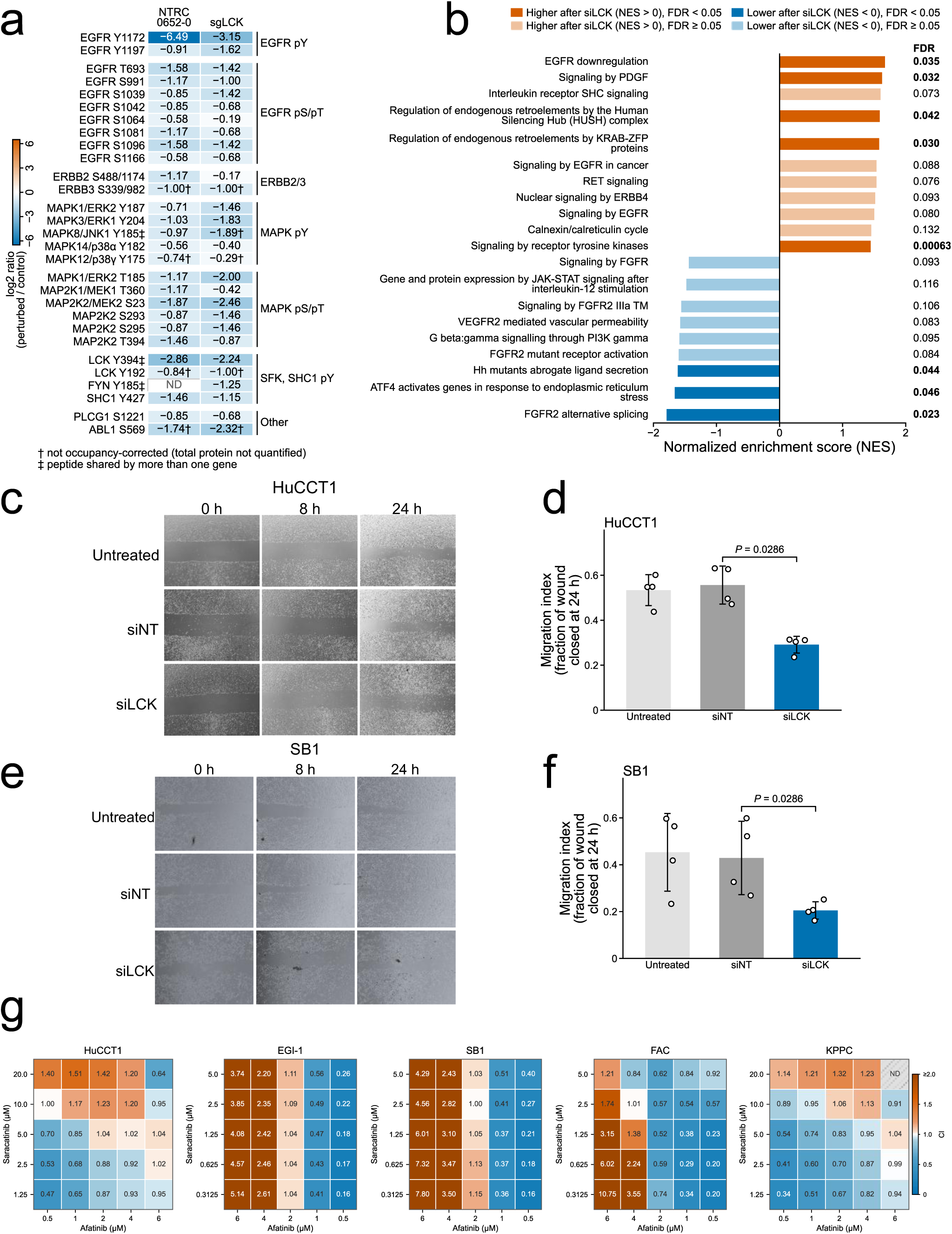
Phosphosite-level and functional consequences of LCK loss, related to Figure 2. (A) SILAC log2 fold changes for EGFR, ERBB2, ERBB3, MAPK and adaptor phosphosites following NTRC 0652-0 treatment or sgLCK disruption in HuCCT1 cells, occupancy-corrected except where marked. Sites are grouped by protein and residue class as indicated. ND, not detected in that arm. †, not occupancy-corrected (total protein not quantified); ‡, peptide shared by more than one gene. (B) Reactome pathway enrichment in siLCK versus siNT HuCCT1 cells for 20 Reactome pathways selected for display, ranked by normalized enrichment score. The Benjamini-Hochberg FDR across the 1,313 sets tested is printed beside each bar; solid bars, FDR < 0.05 (8 of the 20). (C and E) Representative phase-contrast images of scratch wound assays in HuCCT1 (C) and SB1 (E) cells at 0, 8 and 24 h under untreated, siNT and siLCK conditions. No brightness or contrast adjustment was applied to the images. (D and F) Quantification of wound closure in HuCCT1 (D) and SB1 (F) cells, expressed as the fraction of the initial wound area closed at 24 h. Bars show mean ± SD; n = 4 wells per condition from one experiment. P values by exact two-sided Mann-Whitney U test; P = 0.0286 is the smallest value attainable at 4 versus 4. (G) Combination matrices for afatinib and saracatinib across five CCA cell lines. Values are Chou-Talalay combination indices (CI); CI below 1 indicates synergy, and the color scale is shown at right. ND, not determined. Each matrix is one experiment with 3 wells per dose combination, and CI values are single point estimates.

**Figure S3.**
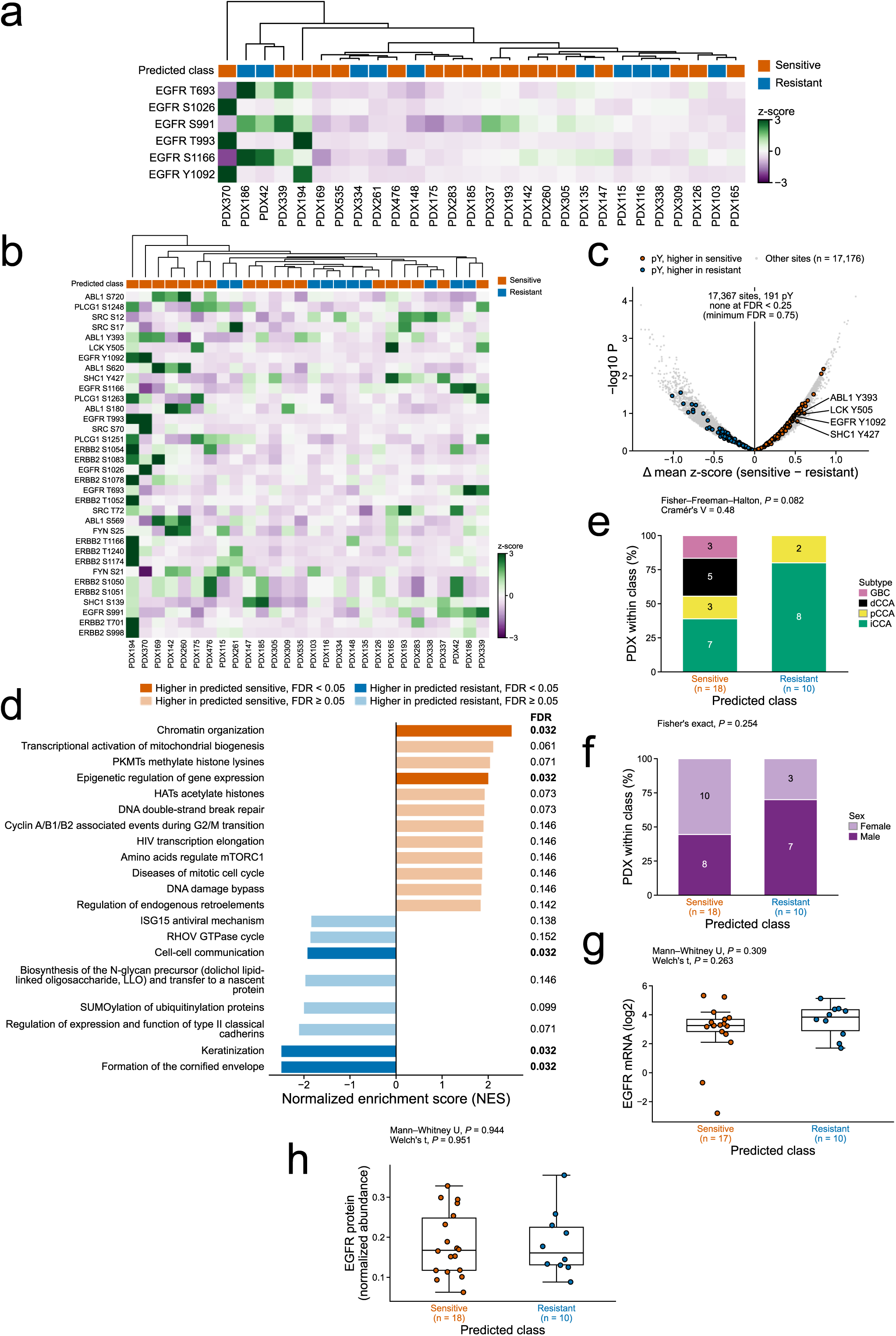
Phosphosite-level characterization of the PDX panel, related to Figure 3. (A) Heatmap of the EGFR phosphosites quantified across 28 PDX models (T693, S991, T993, S1026, S1166 and Y1092), expressed as Z-scores. Column annotation indicates predicted class. Individual EGFR phosphosites did not separate the predicted classes. (B) Heatmap of EGFR, ERBB2, Src-family kinase and associated signaling-protein phosphosites across the same models, expressed as Z-scores. Column annotation indicates predicted class. Columns are clustered independently of (A). (C) Differential phosphosite abundance between predicted-sensitive and predicted-resistant PDX models, as the difference in mean Z-score (sensitive minus resistant) by Welch’s t-test with Benjamini-Hochberg correction. Of 17,367 quantified sites, 191 were tyrosine sites (highlighted); no site reached FDR < 0.05. (D) Reactome pathway enrichment between predicted classes (fgsea, 10,000 permutations, seed 42) on genes ranked by the pooled-variance two-sample t statistic, predicted sensitive (n = 18) versus predicted resistant (n = 10). Of 749 pathways tested, the 12 with the largest positive and the 8 with the largest negative normalized enrichment scores are shown, with FDR printed beside each bar. Solid bars, FDR < 0.05 (5 bars); a sixth pathway at FDR < 0.05, RHO GTPase cycle (FDR 0.046), lies outside the 20 shown. Positive normalized enrichment scores indicate enrichment in predicted-sensitive tumors. (E-H) Sensitive (n = 18) versus resistant (n = 10) class distribution with respect to (E) anatomic subtype (Fisher-Freeman-Halton exact test, P = 0.082; Cramér’s V = 0.48; 6 of 8 cells with expected counts below 5), (F) sex (Fisher’s exact test), (G) EGFR mRNA (n = 27 with PDX142 excluded: 17 sensitive and 10 resistant; Mann-Whitney U test P = 0.3093; Welch’s t-test P = 0.2626; high is at or above the median) and (H) EGFR protein (Mann-Whitney U test and Welch’s t-test as printed in the panel).

**Table S1. Proteins identified by proximity labeling and mean normalized counts of T-cell genes and EGFR in HuCCT1 cells, related to Figure 1**. Table S1A lists the 116 proteins recovered by anti-biotin (BioSITe) enrichment and mass spectrometry from biotin-treated LCK-TurboID and control cells, ranked by proximity enrichment (log2 with a pseudocount of 155,000), with the arm or arms in which each protein was detected (LCK-TurboID only, both arms or control only), the number of biotinylated peptides and, where measured, the EGF/control and afatinib/control ratios. Table S1B lists the 93 proteins quantified in the EGF and afatinib experiment (Figure 1H), with the label-free intensity of each protein in biotin-treated LCK-TurboID cells alone, with EGF and with afatinib, the number of biotinylated peptides and the EGF/control, afatinib/control and afatinib/EGF ratios. Table S1C lists the mean normalized count of CD247 (CD3ζ), CD3D, ZAP70, LAT and EGFR in HuCCT1 RNA-sequencing data, with the number of samples and the definition of the value. Mean normalized count is the DESeq2 baseMean, the mean of median-of-ratios size-factor-normalized counts across all six HuCCT1 RNA-seq samples, with genes of total count below 10 removed before size-factor estimation.

**Table S2. Clinicopathological and molecular characteristics of the patient-derived xenograft models, related to Figure 3**.

**Table S3. EGFR classifier outputs for the PDX models and the human cohorts, related to Figures 3 and 4**. The sheet PDX_28_models lists, for each of the 28 PDX models, the EGFR KEA3 rank and predicted class. The sheets Fudan_214 and Mayo_43 list, for each tumor in the Fudan (n = 214) and Mayo (n = 43) cohorts, the standardized EGFR activity score, the predicted class and the rank of the tumor by score within its cohort (rank 1 is the highest score).

**Table S4. Supporting analyses for the EGFR activity classifier and the LCK-substrate association, related to Figures 3 and 4**. The sheet T-cell adjustment lists, for the Fudan (n = 214 tumors, 208 with RNA-seq) and Mayo (n = 43) cohorts, the Spearman correlation between the LCK-substrate score and the EGFR classifier score unadjusted and after adjustment for a T-cell score, with the 95% confidence interval, P value and n of each, and the correlation of the T-cell score with each of the two scores; the covariate is the mean z-score of six genes in the Fudan cohort and eight in the Mayo cohort, and alternative covariates are listed. The sheet Survival by class lists, for the Fudan (n = 207) and Mayo (n = 40) cohorts, the patients and deaths in each predicted class, the log-rank P, the hazard ratio of predicted sensitive versus predicted resistant with its 95% confidence interval, the hazard ratio per standard deviation of the EGFR activity score and the Kaplan-Meier median overall survival. The sheet Single-feature proxies lists, for the six treated PDX models, the value, the sensitive or resistant call and the concordance with the observed afatinib response class for EGFR protein, batch-corrected EGFR pY1092, raw EGFR pY1092, a curated EGFR-substrate count and the EGFR KEA3 rank class, with a summary row for each proxy.

## METHODS

### Cell culture

Human CCA cell lines HuCCT1, RBE and EGI-1 and murine CCA cell lines SB1, FAC and KPPC (KRAS^G12D^/TP53^−/−^) were maintained in DMEM supplemented with 10% fetal bovine serum and 1% Primocin at 37 °C in a humidified atmosphere containing 5% CO₂. SB1 and FAC cells were established in-house as described previously.^39,40^ Human cell lines were authenticated by short tandem repeat profiling. All cell lines were tested for mycoplasma contamination using MycoStrip (Invivogen, cat. no NC2683201) and were used within 15 passages.

### Generation of doxycycline-inducible HuCCT1-sgLCK cells

HuCCT1-sgLCK cells were generated as described previously.^10^ Briefly, an sgRNA targeting LCK (5′-TCCCGACCCACTGGTTACCTACGA-3′) was introduced by lentiviral transduction into HuCCT1 together with FUCas9Cherry plasmid. Before experiments, lentivirally transduced HuCCT1-sgLCK cells were cultured for three passages in medium containing 7.5 µg/ml doxycycline hyclate (Sigma-Aldrich, cat. no. D9891-1G). Loss of LCK protein was confirmed by immunoblotting.Generation of HuCCT1-sgLCK/TurboID-LCK-HA cells

An LCK-TurboID-HA fusion construct under the control of the EF-1α promoter (pEF6-Lck-TurboID; Addgene, cat. no. 159433)^41^ was introduced into doxycycline-inducible HuCCT1-sgLCK cells using Lipofectamine 3000 (Invitrogen, cat. no. L3000001) according to the manufacturer’s instructions. Stable cells were selected with blasticidin (Gibco, cat. no. A1113903) at 10 µg/ml for 14 days to generate HuCCT1-sgLCK/LCK-TurboID-HA cells. Expression of the fusion protein was confirmed by immunoblotting for the HA tag. Parental HuCCT1 cells were maintained in parallel as controls. The RBE LCK-TurboID-HA line used for the co-immunoprecipitation in Figure 1F was generated in this study by transfecting the same construct with the same transfection and selection procedure.

### siRNA transfection

HuCCT1 and SB1 cells were transfected with an siRNA targeting LCK (siLCK; Dharmacon, cat no. L-003151-00-0010 and L-043878-00-0010) or a non-targeting control siRNA (siNT; Dharmacon cat no. D-001810-01-20) at a final concentration of 12.5 nM using Lipofectamine RNAiMax (Invitrogen, cat no. 13778030), according to the manufacturer’s instructions. Downstream assays were performed 48-72h after transfection.

### Reagents and inhibitors

Afatinib (MedChemExpress, cat no. 850140-72-6) saracatinib (MedChemExpress, cat no. 379231-04-6), dasatinib (MedChemExpress, cat no. 302962-49-8) and the selective LCK inhibitor NTRC 0652-0 (kindly provided by Netherlands Translational Research Center B.V.) were dissolved in DMSO and stored at −20 °C. For in vitro experiments, the final DMSO concentration did not exceed 0.1%, and vehicle-treated controls received an equivalent volume of DMSO.

### Cell viability and combination index assays

For half-maximal inhibitory concentration (IC₅₀) measurements, cells were seeded in 96-well plates at 3,000 cells per well in triplicate on day 0. On day 1, cells were treated with serial dilutions of afatinib (10-fold, 0.1 nM to 100 µM) or saracatinib (2-fold, 48.8 nM to 100 µM) in DMSO (0.1%). After 48 h, CellTiter-Glo reagent (Promega, cat. no. G9242) was added according to the manufacturer’s instructions, and luminescence was measured using a BioTek Synergy H1 microplate reader. Dose-response curves and IC₅₀ values were generated in GraphPad Prism v.9.1.0.

For drug-combination studies, afatinib and saracatinib were combined in a 5 × 5 dose matrix with two-fold dilutions, in technical triplicate within a single experiment. The fractional effect (fa) was calculated as fa = g/100, where g denotes the percentage growth inhibition. The combination index (CI) was calculated by the Chou-Talalay method^42^ as CI = D_A/Dx_A + D_B/Dx_B, where D_A and D_B are the doses of afatinib and saracatinib used in combination to achieve a given effect, and Dx_A and Dx_B are the single-agent doses required for the same effect, obtained by inverting four-parameter log-logistic dose-response models. Where a row or column of the matrix had fewer than five evaluable points, a three-parameter log-logistic model (top fixed at 1) was used, with linear interpolation as a fallback. CI values below 1 were considered synergistic, equal to 1 additive and above 1 antagonistic.

### Cell proliferation assay

HuCCT1 cells transfected with siLCK or siNT were seeded in 96 well format at 3000 cells per well. Cell numbers were measured on days 0, 1, 2 and 3 using Celigo Image Cytometer (Revvity). Each condition comprised 6 wells (5 siNT wells at day 0) in one experiment.

### Wound-healing assay

HuCCT1 and SB1 cells were seeded in 6-well plates and grown to confluence. The monolayers were scratched using a P200 pipette tip, washed once with PBS to remove detached cells and imaged at 0, 8 and 24 h. Wound area was quantified using ImageJ. The migration index was calculated as the fraction of the initial wound area closed at 24 h. Each condition comprised 4 wells in one experiment.

### Immunoblotting

Cells were lysed in RIPA buffer supplemented with protease (Roche, cat no. 11873580001) and phosphatase inhibitors (Roche, cat no. 04906837001) for 10 mins on ice. Lysates were clarified by centrifugation at 15 000 rpm for 15 mins at 4 °C, and protein concentrations were determined using a BCA assay (Thermo Scientific, cat no. 23227). Equal amounts of protein (25 µg) were resolved by SDS–PAGE on 10% gels and transferred to PVDF membranes. Membranes were blocked in 5% BSA prepared in TBS containing Tween-20 for 1h at room temperature and incubated overnight at 4 °C with primary antibodies against phospho-EGFR Y1092 (Invitrogen, cat no. MA5-33133 1:1000), total EGFR (Cell Signaling Technology, cat no: 4267S 1:1000), LCK (Cell Signaling Technology, cat no: 2752S 1:1000), HA (Cell Signaling Technology, cat no: 3724S 1:1000) and β-actin (Santa Cruz, cat no. sc-47778, 1:1000) Membranes were washed and incubated with horseradish peroxidase-conjugated secondary antibodies (Invitrogen, cat no:62-6520 and 31460 1:3000) for 1h at room temperature. Signals were detected using ECL chemiluminescence (GE Healthcare). Biotinylated proteins were detected using streptavidin–horseradish peroxidase (Sigma-Aldrich, cat no. S5512)

### Co-immunoprecipitation

HuCCT1-sgLCK/LCK-TurboID-HA and RBE LCK-TurboID-HA cells were maintained under basal culture conditions and lysed in RIPA buffer supplemented with protease and phosphatase inhibitors. Lysates were clarified by centrifugation and precleared with protein A/G beads for 30 mins at 4 °C. For each immunoprecipitation, 1 mg of total protein was incubated overnight at 4 °C with an anti-EGFR antibody, an anti-HA antibody or an equivalent amount of A/G beads. Immune complexes were captured using protein A/G magnetic beads, (Pierce Classic Magnetic IP/Co-IP kit, cat no 88804) overnight at 4 °C. Beads were washed twice with wash buffer, and bound proteins were eluted using Sample Buffer at RT. Input samples corresponded to 5% of the lysate used for immunoprecipitation. Eluates were analyzed by immunoblotting for EGFR and HA as described above.

### In vitro kinase assay

Recombinant EGFR (BPS Bioscience, cat no. 40187) and recombinant LCK (Promega, cat no. V2691) were combined in kinase buffer according to manufacturer protocol in the presence or absence of ATP and poly(4:1 Glu,Tyr) substrate. Where indicated, dasatinib (500 nM) was added, and reactions were preincubated for 15 min at room temperature before addition of ATP. Reactions were incubated at 30 °C for 15 min.

For immunoblot-based analysis, reactions were terminated by adding Laemmli buffer and analyzed by immunoblotting for phospho-EGFR Y1092, LCK and EGFR as described above. For quantitative analysis, kinase activity was measured using the ADP-Glo Kinase Assay (Promega, cat. no V6930) according to the manufacturer’s instructions. Luminescence was recorded using a BioTek plate reader, and data are presented as relative luminescence units from three technical replicates.

### TurboID proximity labeling

HuCCT1-sgLCK/LCK-TurboID-HA and control cells were seeded at 50 000 cells/well and treated with doxycycline (7.5 µg/ml) for 9 days to deplete endogenous LCK before proximity labeling. For EGF/afatinib treatment studies, EGF (100 ng/ml) or afatinib (250 nM) was added into the media 30 minutes before biotin labeling. Biotin was added to the culture medium at a final concentration of 50 µM, and cells were incubated for 30 minutes at 37 °C. Labeling was terminated by removing the biotin-containing medium and washing the cells five times with ice-cold PBS to remove unincorporated biotin. Cells were then harvested for proteomic analysis. Biotinylated peptides were enriched using the BioSITe method described below. The biotin concentration was selected on the basis of a titration of 50, 100 and 200 µM biotin assessed by streptavidin immunoblotting.

### Proteomic analysis of biotinylated peptides

HuCCT1-sgLCK/LCK-TurboID-HA and parental HuCCT1 cells were harvested for proteomic analysis. Proteins were extracted in lysis buffer containing 8 M urea and 50 mM triethylammonium bicarbonate, reduced with 10 mM dithiothreitol and alkylated with 30 mM iodoacetamide. Samples were digested overnight at 37 °C with gentle agitation using TPCK-treated trypsin at an enzyme-to-protein ratio of 1:20. Peptides were desalted using SEP-PAK C18 cartridges.

Biotinylated peptides were enriched using the BioSITe method as described previously.^43^ Briefly, peptides were resuspended in capture buffer containing 50 mM Tris, 150 mM NaCl and 0.5% Triton X-100 and incubated for 2 h at 4 °C with anti-biotin antibody bound to protein G beads. After removal of the supernatant, beads were washed three times with capture buffer, three times with 50 mM Tris and twice with water. Biotinylated peptides were eluted with 80% acetonitrile containing 0.2% trifluoroacetic acid, desalted on a C_18_ reversed-phase column and dried.

Peptides were analyzed on an Orbitrap Exploris 480 mass spectrometer coupled to an Ultimate 3000 liquid chromatography system. Samples were loaded onto a C_18_ trap column (Optima, 2 cm × 100 µm, 100 Å) at 8 µl/min and separated on a C_18_ analytical column (75 µm × 40 cm, 1.7 µm) at 400 nl/min. Mobile phase A consisted of 0.1% formic acid in water, and mobile phase B consisted of 0.1% formic acid in 80% acetonitrile. Peptides were separated using a 30% mobile phase B gradient over 60 min. Data were acquired in data-dependent acquisition mode with a cycle time of 2 sec. MS1 and MS2 spectra were acquired at resolutions of 120,000 and 30,000, respectively. Precursor ions were isolated using a 1.2 m/z isolation window and fragmented at a normalized collision energy of 30.

Raw data were searched against the UniProt human protein database using SEQUEST in Proteome Discoverer v.3.0 (Thermo Fisher Scientific). Searches were performed with trypsin specificity and up to three missed cleavages. Carbamidomethylation of cysteine was specified as a fixed modification, whereas methionine oxidation and lysine biotinylation were specified as variable modifications. Peptide-spectrum matches were filtered to a false-discovery rate of 1% using the Percolator node. Proteins identified in biotin-treated HuCCT1-sgLCK/LCK-TurboID-HA cells were compared with those identified in biotin-treated parental HuCCT1 cells to define the LCK proximity interactome. Proximity enrichment was calculated as log_2_((I_TurboID_ + 155,000)/(I_control_ + 155,000)), where I is the protein intensity and 155,000 is a pseudocount equal to half the smallest non-zero intensity, and proteins were ranked by this value.

### STRING network analysis

Proteins in the LCK proximity interactome were analyzed using STRING v12 with Homo sapiens specified as the reference organism. The input comprised the 83 proteins detected exclusively in fusion-expressing cells. Interactions were retained at a minimum confidence score of 0.7 using experiments, databases, co-expression, text mining, neighborhood, gene fusion and co-occurrence as active evidence channels. The network displayed in Figure 1E comprises the 38 proteins with at least one interaction at this threshold (37 interactions; lowest combined score 0.706); the other 45 proteins are not displayed. Network topology was assessed using protein degree, and networks were visualized in the STRING web interface.

### Post-translational modification signature enrichment analysis

EGFR residues follow UniProt P00533 numbering. The tyrosine phosphoproteomic dataset analyzed in Figure 2 was generated previously by three-state SILAC of HuCCT1 cells treated with NTRC 0652-0 (1 µM, 24 h) or after genetic deletion of LCK (HuCCT1-sgLCK cells, doxycycline induction for 72 h).^11^ Light (control), medium (NTRC 0652-0) and heavy (sgLCK) cultures were lysed and pooled for one three-state experiment, and the data are available in PRIDE under accession PXD026925. For post-translational modification signature enrichment analysis (PTM-SEA), phosphosite scores were the log_2_ of the raw perturbed-to-control ratio, with zero values set to 0.001 before transformation, from the NTRC 0652-0 and sgLCK arms. The PTM-SEA input comprised the 144 tyrosine phosphosite rows and 9,161 serine and threonine phosphosite rows from IMAC enrichment, with multi-site rows expanded to one entry per site. PTM-SEA was run with ssGSEA2.0 against PTMsigDB v2.0.0 using a weight of 0.75, area under the running enrichment score as the statistic, Z-score correlation, a minimum overlap of 3 sites and 1,000 permutations.^28^ Signatures with a false-discovery rate of < 0.05 within each perturbation arm were considered significant.

### RNA sequencing and differential expression analysis

Total RNA was isolated with TRIzol Reagent (Invitrogen, cat no. 15596026) at the 72h timepoint from HuCCT1 cells transfected with siLCK or siNT according to the manufacturer’s instructions. Library preparation and RNA sequencing was performed by Plasmidsaurus. Differential expression analysis was performed using DESeq2 v1.46.0 with a minimum count filter of 10. Genes with false-discovery rate <0.05 were considered differentially expressed.^44^

### Gene-set enrichment analysis

Genes were ranked by the DESeq2 (v1.46.0) Wald statistic from the siLCK versus siNT comparison and analyzed by gene-set enrichment analysis using the fgsea R package v1.32.4. Gene sets were obtained from msigdbr v26.1.1 (MSigDB 2026.1.Hs, human) and comprised the Hallmark, KEGG MEDICUS and Reactome (C2 CP) collections. Gene sets containing 15 to 500 genes were tested (1,313 sets), using nPermSimple = 10000 permutations and seed 42. P values were adjusted by the Benjamini–Hochberg method across all tested gene sets, and sets with FDR < 0.05 were considered significantly enriched (346 sets). Normalized enrichment scores and FDR values are reported for each gene set shown.

### DepMap dependency and co-dependency analyses

Chronos gene-effect scores and cell-line annotations were obtained from the DepMap portal (release 26Q1).^45^ Biliary tract cell lines were defined by the DepMap lineage annotation (n = 34). Co-dependency between LCK and EGFR was assessed by Spearman correlation of CRISPR gene-effect scores across these lines, and the same correlation with EGFR was calculated for each of 418 kinases. P values were not adjusted for the number of kinases examined.

### Patient-derived xenograft generation and treatment

Patient-derived xenografts (PDXs) from resected biliary tract cancers were generated at Mayo Clinic under an institutional review board-approved protocol (IRB 19-012104). Tumor implantation was performed as described previously.^23^ Briefly, resected tumors were implanted bilaterally into 6-8-week-old NOD/SCID mice. After establishment of each PDX model, 4-mm³ tumor fragments were reimplanted for treatment studies. Mice received afatinib (30 mg/kg) or methylcellulose/Tween-80 vehicle by oral gavage once daily after tumors were established (day 1 of treatment). Tumor dimensions were measured twice weekly from the first day of treatment using calipers, and tumor volume was calculated as length × width². Change in tumor volume was calculated as the final volume minus the baseline volume. Dynamic growth rate was calculated for each mouse as the difference between its tumor volume on the last recorded day and on day 1, divided by the number of days between the two measurements (13 days for day 14, 12 days for day 13, and 7 days for the one censored PDX165 vehicle mouse), in mm³/day. A total of 4-5 mice were included per treatment group for each PDX model. PDX experiments used mice of both sexes, and no difference between sexes was observed previously. Syngeneic experiments used male C57BL/6 mice. Investigators measuring tumors were blinded to treatment allocation.

### Syngeneic orthotopic murine CCA models and treatment

All animal experiments were approved by the Institutional Animal Care and Use Committee before study initiation. Murine SB1 (myr-AKT/YAP^S127A^) and FAC (FBXW7/Akt) cells were generated in-house as described previously, and orthotopic CCA models were established as described previously.^39^ Briefly, SB1 or FAC cells were harvested and resuspended in a 1:1 mixture of Matrigel (Thermo Fisher Scientific, cat. no. CB40234) and DMEM (Thermo Fisher Scientific, cat. no. 10566016). Six-week-old male C57BL/6 mice were anesthetized with isoflurane, and 1 × 10 SB1 or FAC cells in 40 µl of the Matrigel–DMEM mixture were injected into the left hepatic lobe using a 27-gauge needle.

Mice were randomized to receive afatinib (30 mg/kg) or methylcellulose/Tween-80 vehicle by oral gavage once daily. Treatment was initiated 2 weeks after implantation of SB1 cells or 3 weeks after implantation of FAC cells and continued for 14 days. At the humane endpoint, mice were euthanized according to the approved IACUC protocol, and liver, tumor and blood samples were collected for downstream analyses. Tumors were weighed at necropsy.

### Estimation of drug sensitivity from phosphoproteomic profiles

Kinase activity ranks for the 28 PDX models were obtained from kinase enrichment analysis of the phosphosites enriched in each model. Total proteome and phosphoproteome (pSTY and pY) intensities were each normalized for differences in total signal between samples, and each phosphosite signal was then divided by the signal of the corresponding total protein. Batches were merged on features present in all samples and corrected with ComBat^46,47^ (R package, default settings); outlier samples that remained divergent after correction were removed and the correction was repeated. For each model, phosphosites with a signal more than 4-fold above the median of the cohort were designated enriched. The enriched phosphosites of each model were submitted to KEA3^31^ through the KEA3web query interface in 2021; a version number was not recorded. KEA3 integrates multiple kinase-substrate interaction resources and returns a ranked list of predicted kinases for each model. The integrated MeanRank output was used, with rank 1 representing the strongest inferred kinase activity, and the EGFR rank was extracted for each model, with a low rank indicating greater inferred EGFR kinase activity. Models with an EGFR rank below 10 were classified as predicted sensitive, and models with an EGFR rank of 10 or higher were classified as predicted resistant. Per-model ranks and classes are provided in Table S3, so that the classes can be verified without repeating the web query. For the single-feature proxy comparison (Table S4), a model was called sensitive when its proxy value (EGFR protein, batch-corrected EGFR pY1092 z-score, raw EGFR pY1092 intensity, or the number of the 41 curated EGFR-substrate genes in the KEA3 input list of the model) was above the median of that proxy over the 22 untreated models (18 for raw EGFR pY1092, because four untreated models lack the raw intensity), and the call was compared with the observed afatinib response class of each of the six treated models; the EGFR-rank class was called by EGFR rank below 10.

In the Fudan and Mayo cohorts, the analogous cohort-standardized score was used (see below).

### Kinase signature enrichment between predicted classes in PDX models

Batch-corrected phosphosite Z-scores (17,367 sites) from 28 PDX models were compared between the 18 models predicted sensitive and the 10 models predicted resistant by the EGFR activity classifier. A Welch two-sample t-test (sensitive versus resistant) was applied at each site with Benjamini-Hochberg correction. No individual site reached a false-discovery rate of <0.05 or <0.25, so the signed t statistics, positive when higher in predicted-sensitive models, were used as the ranked list. Post-translational modification signature enrichment analysis (PTM-SEA) was performed with ssGSEA2.0 against PTMsigDB v2.0.0 using a weight of 0.75, area under the running enrichment score, rank-based correlation, 1,000 permutations, a minimum overlap of 3 sites and a random seed of 42. Normalized enrichment scores (NES) and false-discovery rates are reported, with positive NES indicating enrichment in predicted-sensitive models and negative NES indicating enrichment in predicted-resistant models. In Figure 3J, all KINASE-PSP (PhosphoSitePlus) and KINASE-iKiP signatures with a false-discovery rate of <0.25 in either direction are shown, together with the LCK, SRC, ABL1 and ERBB2 signatures regardless of false-discovery rate.^48,49^ When a kinase had both a PSP and an iKiP signature in the same direction, the signature with the larger absolute NES was kept. When the directions differed, both were kept and labeled by library. Dot color indicates NES and dot size indicates-log10 false-discovery rate.

### Class comparison in the PDX panel

Batch-corrected phosphosite Z-scores (17,367 sites) were compared between the 18 predicted-sensitive and 10 predicted-resistant PDX models by Welch’s two-sample t-test at each site, with Benjamini-Hochberg correction; 191 tyrosine sites are highlighted (Figure S3C). Pathway enrichment of the phosphoproteome was performed as described for the Fudan cohort against the Reactome collection (Figure S3D). Application of the EGFR activity classifier to human cohorts

#### Application of the EGFR activity classifier to human cohorts

The EGFR activity score was computed in two independent cholangiocarcinoma cohorts without refitting: the Fudan FU-iCCA cohort (n = 214) and a Mayo Clinic CCA cohort (n = 43 CCA tumors). In each cohort, phosphosite intensities were converted to Z-scores across the tumors, phosphosites whose Z-score in a given tumor exceeded 1.96 were submitted to KEA3 as described above, and the resulting EGFR activity score was standardized as a Z-score within the cohort, because the cohorts were profiled on different platforms and the PDX rule is an absolute KEA3 rank (below 10). This site-selection criterion differs from the one used for the PDX models (sites more than 4-fold above the cohort median of batch-corrected, protein-normalized phosphosite signal), so the cohort scores are analogous to, and not identical with, the PDX ranks. Tumors with a standardized score of 0 or higher were classified as predicted sensitive, and those below 0 as predicted resistant. The date of the KEA3web queries for the cohort scores was not recorded. Class sizes follow from the standardization and are not calibrated to drug response, because the threshold is relative to each cohort. Per-tumor scores and classes are provided in Table S3.

### LCK kinase-activity scoring in human cohorts

LCK-substrate phosphorylation scores were computed from the published log2-ratio phosphosite tables of each cohort (Fudan, Table S1E of Dong et al.;^22^ Mayo, Table S6 of Mun et al., TMT ratios^23^). Phosphosites were matched at the gene level, regardless of residue, to the LCK substrate genes in OmniPath v3.14.0^50^ and the score of each sample was the unweighted mean of its observed matched sites; the decoupleR run_wmean function applied per sample to the observed sites gives the same values.^50^ The Fudan correlation in Figure 4C uses all 214 classified tumors. Of the matched sites, 182 of 196 (Fudan) and 249 of 279 (Mayo) are serine or threonine sites, so the score reports LCK-substrate phosphorylation and not a tyrosine readout. Scores were recomputed from the published tables with kinase-substrate relationships from an OmniPath snapshot downloaded on 24 March 2026. Spearman P values are two-sided and asymptotic, confidence intervals for the correlations were obtained by the Fisher z transformation, the Fudan and Mayo correlations were compared by the Fisher z test for independent correlations, and the minimum detectable correlation was calculated for 80% power at a two-sided α of 0.05. Partial Spearman correlations, adjusting for a T-cell covariate, were calculated as the correlation of rank residuals, with P values from the t distribution and 95% confidence intervals by the Fisher z transformation. The covariate was the mean z-score of CD3D, CD3E, CD3G, CD2, CD8A and CD4 in the Fudan RNA-seq data, whose table has no TRAC or TRBC1 rows, and of these six genes plus TRAC and TRBC1 in the Mayo RNA-seq data (the six-gene version gives the same result in the Mayo cohort); LCK was excluded from the covariate (Table S4).

### Mutation frequency analysis

Somatic mutation calls for each cohort were obtained from the published datasets for each cohort.^22,23^ For each gene, mutation frequencies were compared between predicted-sensitive and predicted-resistant tumors using two sided Fisher’s exact test. P values were adjusted for multiple comparisons using the Benjamini–Hochberg method.

### Differential phosphosite and gene-set enrichment analysis

Of the 18,347 phosphosites, 14,935 sites with no more than 40% missing values across the 214 classified tumors were retained for differential analysis, and remaining missing values were imputed by the row mean. Differential analysis was performed with limma v3.62.2 on the Z-scored intensities.^51^ A linear model with a two-group design (predicted resistant as the reference) was fit with lmFit, followed by empirical Bayes moderation with eBayes, and the contrast tested predicted sensitive versus predicted resistant. P values were adjusted by the Benjamini-Hochberg method, and 1,146 sites were higher in predicted-sensitive tumors and 1,214 were higher in predicted-resistant tumors at FDR < 0.05 (Figure 4F). For pathway analysis, phosphosites were collapsed to the gene level by retaining, for each gene, the site with the largest absolute limma t statistic, and genes were ranked by this signed statistic. Preranked gene-set enrichment analysis was performed with fgsea v1.32.4 using 10,000 permutations and bidirectional scoring against the Reactome (C2 CP; 887 sets) and KEGG Legacy (C2 CP; 163 sets) collections from MSigDB 2026.1.Hs accessed through msigdbr v26.1.0, with set sizes of 10 to 500 genes.^52^ P values were adjusted by the Benjamini-Hochberg method, and normalized enrichment scores and FDR values are reported for the pathways shown in Figure 4G. Analyses were performed in R v4.4.3.

### Survival analysis

Overall survival was compared between tumors with high and low EGFR expression, defined by a median split of EGFR mRNA abundance within each cohort. Survival distributions were estimated using the Kaplan–Meier method and compared using the log-rank test.

### Statistical analysis

The statistics for the ADP-Glo kinase assay, the siLCK growth curve, the syngeneic tumor weights, the PDX treatment and response comparisons, the mutation-frequency comparison between predicted classes and the Spearman correlations and their comparison between the human cohorts were computed in Python v3.11.15 (SciPy v1.17.1). The wound-closure comparison was computed in Python and the overall survival comparisons in R with the survival package; the versions used for these two were not recorded. The class-association tests for the 28 PDX models and the Fudan and Mayo cohorts were computed in Python v3.13.14 (SciPy v1.18.0), with R v4.4.3 (fisher.test) for the exact tests. The sequencing, differential phosphosite and enrichment analyses were run in R v4.4.3 and are described in the corresponding sections. All analyses are recorded with the archived analysis outputs. Dose-response curves and IC_50_ values were fitted in GraphPad Prism v.9.1.0. Unless otherwise indicated, data are presented as mean ± s.d. The number of wells, reactions, mice or tumors that n represents is given in the figure legends. ADP-Glo reactions and the siLCK growth curve were compared using two-sided Welch’s t-tests, with growth-curve P values adjusted by the Benjamini-Hochberg method across days. For each PDX model, final tumor-volume changes were compared between treatment groups using a two-sided Student’s t-test, and dynamic growth rates were compared using a two-sided exact Mann-Whitney U test. P values across the six PDX models were not adjusted. Tumor weights in the syngeneic orthotopic models and wound-closure fractions were compared using exact two-sided Mann-Whitney U tests. Concordance between predicted class and observed afatinib response was assessed using a one-sided Fisher’s exact test. Correlations are reported as Spearman rank-correlation coefficients; for the six treated PDX models, P values were calculated by exact permutation (all 720 rank orderings). Associations between predicted class and clinical or molecular variables were tested by two-sided Mann-Whitney U tests for continuous variables and two-sided Fisher’s exact tests for categorical variables, with the Fisher-Freeman-Halton exact test for anatomic subtype in the PDX panel. P values from omics analyses were adjusted by the Benjamini-Hochberg method. P < 0.05 was considered statistically significant.

## REFERENCES

1 Banales, J. M. et al. Cholangiocarcinoma 2026: status quo, unmet needs and priorities. Nat Rev Gastroenterol Hepatol 23, 65–96 (2026). 10.1038/s41575-025-01153-w

2 Ilyas, S. I., Khan, S. A., Hallemeier, C. L., Kelley, R. K. & Gores, G. J. Cholangiocarcinoma — evolving concepts and therapeutic strategies. Nature Reviews Clinical Oncology 15, 95–111 (2018). 10.1038/nrclinonc.2017.157

3 Kelley, R. K. et al. Pembrolizumab in combination with gemcitabine and cisplatin compared with gemcitabine and cisplatin alone for patients with advanced biliary tract cancer (KEYNOTE-966): a randomised, double-blind, placebo-controlled, phase 3 trial. Lancet 401, 1853–1865 (2023). 10.1016/S0140-6736(23)00727-4

4 Casak, S. J. et al. FDA Approval Summary: Durvalumab and Pembrolizumab, Immune Checkpoint Inhibitors for the Treatment of Biliary Tract Cancer. Clin Cancer Res 30, 3371–3377 (2024). 10.1158/1078-0432.CCR-24-0517

5 Bommhardt, U., Schraven, B. & Simeoni, L. Beyond TCR Signaling: Emerging Functions of Lck in Cancer and Immunotherapy. International Journal of Molecular Sciences 20, 3500 (2019).

6 Weiße, J. et al. Identification of lymphocyte cell-specific protein-tyrosine kinase (LCK) as a driver for invasion and migration of oral cancer by tumor heterogeneity exploitation. Molecular Cancer 20, 88 (2021). 10.1186/s12943-021-01384-w

7 Rupniewska, E. et al. Targeting autophagy sensitises lung cancer cells to Src family kinase inhibitors. Oncotarget 9, 27346–27362 (2018). 10.18632/oncotarget.25213

8 Dey, G. et al. LCK facilitates DNA damage repair by stabilizing RAD51 and BRCA1 in the nucleus of chemoresistant ovarian cancer. Journal of Ovarian Research 16, 122 (2023). 10.1186/s13048-023-01194-2

9 Saygin, C. et al. CD55 regulates self-renewal and cisplatin resistance in endometrioid tumors. J Exp Med 214, 2715–2732 (2017). 10.1084/jem.20170438

10 Sugihara, T. et al. YAP Tyrosine Phosphorylation and Nuclear Localization in Cholangiocarcinoma Cells Are Regulated by LCK and Independent of LATS Activity. Mol Cancer Res 16, 1556–1567 (2018). 10.1158/1541-7786.MCR-18-0158

11 Conboy, C. B. et al. LCK inhibition downregulates YAP activity and is therapeutic in patient-derived models of cholangiocarcinoma. Journal of Hepatology 78, 142–152 (2023). 10.1016/j.jhep.2022.09.014

12 Yu, M. et al. Aptamer-directed, nanovesicle-mediated targeting of undruggable molecules in preclinical cholangiocarcinoma models. Hepatology (2025). 10.1097/hep.0000000000001256

13 Blake, S., Hughes, T. P., Mayrhofer, G. & Lyons, A. B. The Src/ABL kinase inhibitor dasatinib (BMS-354825) inhibits function of normal human T-lymphocytes in vitro. Clin Immunol 127, 330–339 (2008). 10.1016/j.clim.2008.02.006

14 Schade, A. E. et al. Dasatinib, a small-molecule protein tyrosine kinase inhibitor, inhibits T-cell activation and proliferation. Blood 111, 1366–1377 (2008). 10.1182/blood-2007-04-084814

15 Vijay, V. et al. Generation of a Biliary Tract Cancer Cell Line Atlas Identifies Molecular Subtypes and Therapeutic Targets. Cancer Discov 15, 1858–1882 (2025). 10.1158/2159-8290.CD-24-1383

16 Fouassier, L. et al. Signalling networks in cholangiocarcinoma: Molecular pathogenesis, targeted therapies and drug resistance. Liver Int 39 **Suppl 1**, 43–62 (2019). 10.1111/liv.14102

17 Wu, Q. et al. EGFR Inhibition Potentiates FGFR Inhibitor Therapy and Overcomes Resistance in FGFR2 Fusion–Positive Cholangiocarcinoma. Cancer Discovery 12, 1378–1395 (2022). 10.1158/2159-8290.Cd-21-1168

18 Jin, H. et al. EGFR activation limits the response of liver cancer to lenvatinib. Nature 595, 730–734 (2021). 10.1038/s41586-021-03741-7

19 Vogel, A. et al. PICCA study: panitumumab in combination with cisplatin/gemcitabine chemotherapy in KRAS wild-type patients with biliary cancer-a randomised biomarker-driven clinical phase II AIO study. Eur J Cancer 92, 11–19 (2018). 10.1016/j.ejca.2017.12.028

20 Malka, D. et al. Gemcitabine and oxaliplatin with or without cetuximab in advanced biliary-tract cancer (BINGO): a randomised, open-label, non-comparative phase 2 trial. Lancet Oncol 15, 819–828 (2014). 10.1016/S1470-2045(14)70212-8

21 Harle, A. et al. Expression of pEGFR and pAKT as response-predictive biomarkers for RAS wild-type patients to anti-EGFR monoclonal antibodies in metastatic colorectal cancers. Br J Cancer 113, 680–685 (2015). 10.1038/bjc.2015.250

22 Dong, L. et al. Proteogenomic characterization identifies clinically relevant subgroups of intrahepatic cholangiocarcinoma. Cancer Cell 40, 70–87.e15 (2022). 10.1016/j.ccell.2021.12.006

23 Mun, D. G. et al. Multiomics combined with machine learning defines unique molecular subtypes of cholangiocarcinoma and identifies TNK1 as a therapeutic target. Hepatology (2025). 10.1097/HEP.0000000000001535

24 Branon, T. C. et al. Efficient proximity labeling in living cells and organisms with TurboID. Nature Biotechnology 36, 880–887 (2018). 10.1038/nbt.4201

25 Szklarczyk, D. et al. The STRING database in 2025: protein networks with directionality of regulation. Nucleic Acids Research 53, D730–D737 (2025). 10.1093/nar/gkae1113

26 Arafeh, R., Shibue, T., Dempster, J. M., Hahn, W. C. & Vazquez, F. The present and future of the Cancer Dependency Map. Nature Reviews Cancer 25, 59–73 (2025). 10.1038/s41568-024-00763-x

27 Dempster, J. M. et al. Chronos: a cell population dynamics model of CRISPR experiments that improves inference of gene fitness effects. Genome Biology 22, 343 (2021). 10.1186/s13059-021-02540-7

28 Krug, K. et al. A Curated Resource for Phosphosite-specific Signature Analysis. Mol Cell Proteomics 18, 576–593 (2019). 10.1074/mcp.TIR118.000943

29 Liberzon, A. et al. The Molecular Signatures Database (MSigDB) hallmark gene set collection. Cell Syst 1, 417–425 (2015). 10.1016/j.cels.2015.12.004

30 Buffière, A. et al. Saracatinib impairs maintenance of human T-ALL by targeting the LCK tyrosine kinase in cells displaying high level of lipid rafts. Leukemia 32, 2062–2065 (2018). 10.1038/s41375-018-0081-5

31 Kuleshov, M. V. et al. KEA3: improved kinase enrichment analysis via data integration. Nucleic Acids Res 49, W304–w316 (2021). 10.1093/nar/gkab359

32 Biscardi, J. S. et al. c-Src-mediated phosphorylation of the epidermal growth factor receptor on Tyr845 and Tyr1101 is associated with modulation of receptor function. J Biol Chem 274, 8335–8343 (1999). 10.1074/jbc.274.12.8335

33 Reinehr, R., Becker, S., Höngen, A. & Häussinger, D. The Src family kinase Yes triggers hyperosmotic activation of the epidermal growth factor receptor and CD95. Vol. 279 (2004) 23977–23987. Journal of Biological Chemistry 279, 31990 (2004). 10.1016/S0021-9258(18)36265-3

34 Modjtahedi, H., Cho, B. C., Michel, M. C. & Solca, F. A comprehensive review of the preclinical efficacy profile of the ErbB family blocker afatinib in cancer. Naunyn Schmiedebergs Arch Pharmacol 387, 505–521 (2014). 10.1007/s00210-014-0967-3

35 Lee, J. et al. Gemcitabine and oxaliplatin with or without erlotinib in advanced biliary-tract cancer: a multicentre, open-label, randomised, phase 3 study. Lancet Oncol 13, 181–188 (2012). 10.1016/s1470-2045(11)70301-1

36 Leone, F. et al. Panitumumab in combination with gemcitabine and oxaliplatin does not prolong survival in wild-type KRAS advanced biliary tract cancer: A randomized phase 2 trial (Vecti-BIL study). Cancer 122, 574–581 (2016). 10.1002/cncr.29778

37 Moehler, M. et al. Safety and efficacy of afatinib as add-on to standard therapy of gemcitabine/cisplatin in chemotherapy-naive patients with advanced biliary tract cancer: an open-label, phase I trial with an extensive biomarker program. BMC Cancer 19, 55 (2019). 10.1186/s12885-018-5223-7

38 Schneider, A. et al. Prospective pan-cancer phosphoproteomics at clinical scale extends therapeutic options in precision oncology. bioRxiv, 2026.2007.2008.737171 (2026). 10.64898/2026.07.08.737171

39 Tomlinson, J. L. et al. Syngeneic murine models with distinct immune microenvironments represent subsets of human intrahepatic cholangiocarcinoma. J Hepatol 80, 892–903 (2024). 10.1016/j.jhep.2024.02.008

40 I Ilyas, S., et al. YAP-associated chromosomal instability and cholangiocarcinoma in mice. Oncotarget 9 (2017).

41 Chua, X. Y., Aballo, T., Elnemer, W., Tran, M. & Salomon, A. Quantitative Interactomics of Lck-TurboID in Living Human T Cells Unveils T Cell Receptor Stimulation-Induced Proximal Lck Interactors. J Proteome Res 20, 715–726 (2021). 10.1021/acs.jproteome.0c00616

42 Chou, T.-C. Drug Combination Studies and Their Synergy Quantification Using the Chou-Talalay Method. Cancer Research 70, 440–446 (2010). 10.1158/0008-5472.Can-09-1947

43 Kim, D. I. et al. BioSITe: A Method for Direct Detection and Quantitation of Site-Specific Biotinylation. J Proteome Res 17, 759–769 (2018). 10.1021/acs.jproteome.7b00775

44 Love, M. I., Huber, W. & Anders, S. Moderated estimation of fold change and dispersion for RNA-seq data with DESeq2. Genome Biology 15, 550 (2014). 10.1186/s13059-014-0550-8

45 Tsherniak, A. et al. Defining a Cancer Dependency Map. Cell 170, 564–576.e516 (2017). 10.1016/j.cell.2017.06.010

46 Johnson, W. E., Li, C. & Rabinovic, A. Adjusting batch effects in microarray expression data using empirical Bayes methods. Biostatistics 8, 118–127 (2007). 10.1093/biostatistics/kxj037

47 Leek, J. T., Johnson, W. E., Parker, H. S., Jaffe, A. E. & Storey, J. D. The sva package for removing batch effects and other unwanted variation in high-throughput experiments. Bioinformatics 28, 882–883 (2012). 10.1093/bioinformatics/bts034

48 Mari, T. et al. In Vitro Kinase-to-Phosphosite Database (iKiP-DB) Predicts Kinase Activity in Phosphoproteomic Datasets. Journal of Proteome Research 21, 1575–1587 (2022). 10.1021/acs.jproteome.2c00198

49 Hornbeck, P. V. et al. PhosphoSitePlus: a comprehensive resource for investigating the structure and function of experimentally determined post-translational modifications in man and mouse. Nucleic Acids Research 40, D261–D270 (2012). 10.1093/nar/gkr1122

50 Türei, D. et al. OmniPath: integrated knowledgebase for multi-omics analysis. Nucleic Acids Research 54, D652–D660 (2026). 10.1093/nar/gkaf1126

51 Ritchie, M. E. et al. limma powers differential expression analyses for RNA-sequencing and microarray studies. Nucleic Acids Research 43, e47–e47 (2015). 10.1093/nar/gkv007

52 Korotkevich, G. et al. Fast gene set enrichment analysis. bioRxiv, 060012 (2021). 10.1101/060012

