## Supplementary Table 2 for "Transactivation of EGFR by LCK underlies an activity-defined dependency that predicts response to EGFR inhibition in cholangiocarcinoma"

**Table S2. Clinicopathological and molecular characteristics of the patient-derived xenograft models, related to Figure 3.**

| **PDX model** | **EGFR KEA3 rank^a^** | **Predicted class^b^** | **Anatomic subtype^c^** | **Sex^d^** | **EGFR protein level^e^** | **EGFR mRNA level^f^** | **Afatinib response^g^** |
| --- | --- | --- | --- | --- | --- | --- | --- |
| PDX147 | 1 | Predicted sensitive | GBC | Female | High | High | Not treated |
| PDX165 | 1 | Predicted sensitive | iCCA | Male | High | Low | Regression |
| PDX169 | 1 | Predicted sensitive | iCCA | Female | Low | Low | Not treated |
| PDX337 | 1 | Predicted sensitive | iCCA | Female | Low | Low | Not treated |
| PDX175 | 2 | Predicted sensitive | dCCA | Female | Low | Low | Not treated |
| PDX476 | 2 | Predicted sensitive | iCCA | Female | Low | Low | Not treated |
| PDX283 | 2 | Predicted sensitive | iCCA | Female | High | High | Not treated |
| PDX142 | 2 | Predicted sensitive | dCCA | Male | High | Excluded | Not treated |
| PDX185 | 3 | Predicted sensitive | dCCA | Male | Low | High | Regression |
| PDX305 | 4 | Predicted sensitive | iCCA | Female | High | High | Not treated |
| PDX194 | 4 | Predicted sensitive | pCCA | Male | High | Low | Regression |
| PDX260 | 4 | Predicted sensitive | pCCA | Male | Low | High | Not treated |
| PDX126 | 5 | Predicted sensitive | GBC | Female | High | High | Not treated |
| PDX193 | 6 | Predicted sensitive | pCCA | Male | High | Low | Not treated |
| PDX535 | 6 | Predicted sensitive | iCCA | Male | Low | Low | Not treated |
| PDX339 | 6 | Predicted sensitive | dCCA | Female | High | High | Not treated |
| PDX370 | 6 | Predicted sensitive | GBC | Female | Low | Low | Not treated |
| PDX309 | 6 | Predicted sensitive | dCCA | Male | Low | Low | Not treated |
| PDX116 | 12 | Predicted resistant | pCCA | Male | High | High | Not treated |
| PDX261 | 14 | Predicted resistant | iCCA | Male | Low | Low | No response |
| PDX135 | 14 | Predicted resistant | iCCA | Male | High | High | No response |
| PDX42 | 14 | Predicted resistant | iCCA | Male | High | High | No response |
| PDX148 | 14 | Predicted resistant | iCCA | Female | Low | High | Not treated |
| PDX115 | 14 | Predicted resistant | iCCA | Male | Low | High | Not treated |
| PDX338 | 16 | Predicted resistant | iCCA | Female | Low | Low | Not treated |
| PDX334 | 29 | Predicted resistant | pCCA | Male | Low | Low | Not treated |
| PDX186 | 37 | Predicted resistant | iCCA | Male | High | High | Not treated |
| PDX103 | 41 | Predicted resistant | iCCA | Female | High | High | Not treated |

Abbreviations: PDX, patient-derived xenograft; EGFR, epidermal growth factor receptor; KEA3, Kinase Enrichment Analysis 3; iCCA, intrahepatic cholangiocarcinoma; pCCA, perihilar cholangiocarcinoma; dCCA, distal cholangiocarcinoma; GBC, gallbladder carcinoma.

^a^ Rank of EGFR among the kinases returned by KEA3 for each model; a lower rank indicates higher inferred EGFR activity. Models are listed in order of EGFR rank.

^b^ Predicted sensitive, EGFR KEA3 rank below 10; predicted resistant, rank of 10 or higher.

^c^ Anatomic subtype as recorded in the institutional PDX registry.

^d^ Sex as recorded in the institutional PDX registry.

^e^ High, EGFR protein abundance at or above the median of the 28 models; low, below the median.

^f^ High, EGFR mRNA level at or above the median of the 27 models with an RNA value; low, below the median. Excluded, the PDX142 RNA value is a constant detection floor and was omitted from RNA analyses.

^g^ Afatinib against vehicle in the six treated models (Figure 3E-G). Regression, tumor volume fell below baseline in every afatinib-treated mouse; no response, tumors continued to grow with no significant difference from vehicle; not treated, the model was not tested in vivo.
